# scMoMaT: Mosaic integration of single cell multi-omics data using matrix tri-factorization

**DOI:** 10.1101/2022.05.17.492336

**Authors:** Ziqi Zhang, Haoran Sun, Ragunathan Mariappan, Xi Chen, Xinyu Chen, Mika S Jain, Mirjana Efremova, Sarah A Teichmann, Vaibhav Rajan, Xiuwei Zhang

## Abstract

Single cell data integration methods aim to integrate cells across data batches and modalities, and obtain a comprehensive view of the cells. Single cell data integration tasks can be categorized into horizontal, vertical, diagonal, and mosaic integration, where mosaic integration is the most general and challenging case with few methods developed. We propose scMoMaT, a method that is able to integrate single cell multi-omics data under the mosaic integration scenario using matrix tri-factorization. During integration, scMoMaT is also able to uncover the cluster specific bio-markers across modalities. These multi-modal bio-markers are used to interpret and annotate the clusters to cell types. Moreover, scMoMaT can integrate cell batches with unequal cell type compositions. Applying scMoMaT to multiple real and simulated datasets demonstrated these features of scMoMaT and showed that scMoMaT has superior performance compared to existing methods. We also show that integrated cell embedding combined with learned bio-markers leads to cell type annotations of higher quality or resolution compared to their original annotations.

## 1 Introduction

The advance in single cell multi-omics technology makes it possible to profile a single cell from multiple modalities. Single cell RNA-sequencing (scRNA-seq) is able to measure the gene expression of individual cells, whereas single cell ATAC-sequencing (scATAC-seq) measures the chromatin accessibility of individual cells. On the other hand, new sequencing technologies have been proposed to profile more than one modality in a cell simultaneously. There exist technologies that are able to profile both protein abundance and gene expression^1^, chromatin accessibility and gene expression^2^, or chromatin accessibility and protein abundance^3^ within a cell at the same time. Integrating cells from multiple modalities provides a comprehensive view of cellular identity and the key features (e.g. chromatin regions, genes, proteins, etc) that define the identity, and can further help to understand the underlying cross-modalities relationships.

Data integration tasks on such single cell data matrices can be categorized into four different scenarios^4^: *horizontal integration*, or termed batch effect removal, refers to the case where all data batches have the same modality. *Vertical integration* refers to the case where a data batch is measured with multiple modalities. *Diagonal integration* refers to the case that neither cells nor modalities are shared between data matrices. *Mosaic integration* is the most general case and can be any combination of horizontal, vertical, and diagonal integration. Considering an *m* × *b* grid that corresponds to *m* modalities and *b* batches, mosaic integration methods aim to integrate any subset of data matrices from this grid.

Various methods have been proposed to deal with these integration scenarios. LIGER^5^ and Seurat v3^6^ were developed for horizontal and diagonal integration tasks. CoupleNMF^7^, UnionCom^8^, MMD-MA^9^, and scDART^10^ were developed for diagonal integration task. Seurat v4^11^, scAI^12^, and MultiVI^13^ were developed for vertical integration task. Recently, new methods have been proposed to work with less restricted integration scenarios. Bridge integration^15^ uses one jointly profiled data batch that includes all modalities as the “bridges” and integrates all data batches using dictionary learning. It requires one batch of cells where all modalities are measured. Recently published methods, MultiMap^14^ and UINMF^16^, can integrate data matrices in mosaic integration scenario, but they both focus on learning cell embedding and do not simultaneously learn a feature embedding along with the marker features of cell types.

Here we propose scMoMaT (**s**ingle **c**ell **M**ulti-**o**mics integration using **Ma**trix **T**rifactorization), a data integration framework that is designed to integrate an arbitrary number of data matrices under mosaic integration scenario (Fig. 1a). Apart from integrating cells, scMoMaT also extracts cell type specific bio-markers across modalities: marker genes (from gene expression modality), marker motifs or regions (from chromatin accessibility modality) and marker proteins (from protein abundance modality), etc. It extracts the bio-marker not only from the original feature of the data matrices, but also from the features that are generated by other methods. For example, users can add motif deviation matrices (learned from the original scATAC-seq matrix through chromVAR^17^, representing the motif activities within cells) to the input, and scMoMaT is able to extract the motif markers in addition to the bio-markers from the original modalities. These bio-markers can be used to annotate cell types with evidence from multiple modalities. In addition, scMoMaT does not assume cells to have similar distribution across batches, which makes scMoMaT capable of integrating cell batches with disproportionate cell type composition.

**Figure 1.**
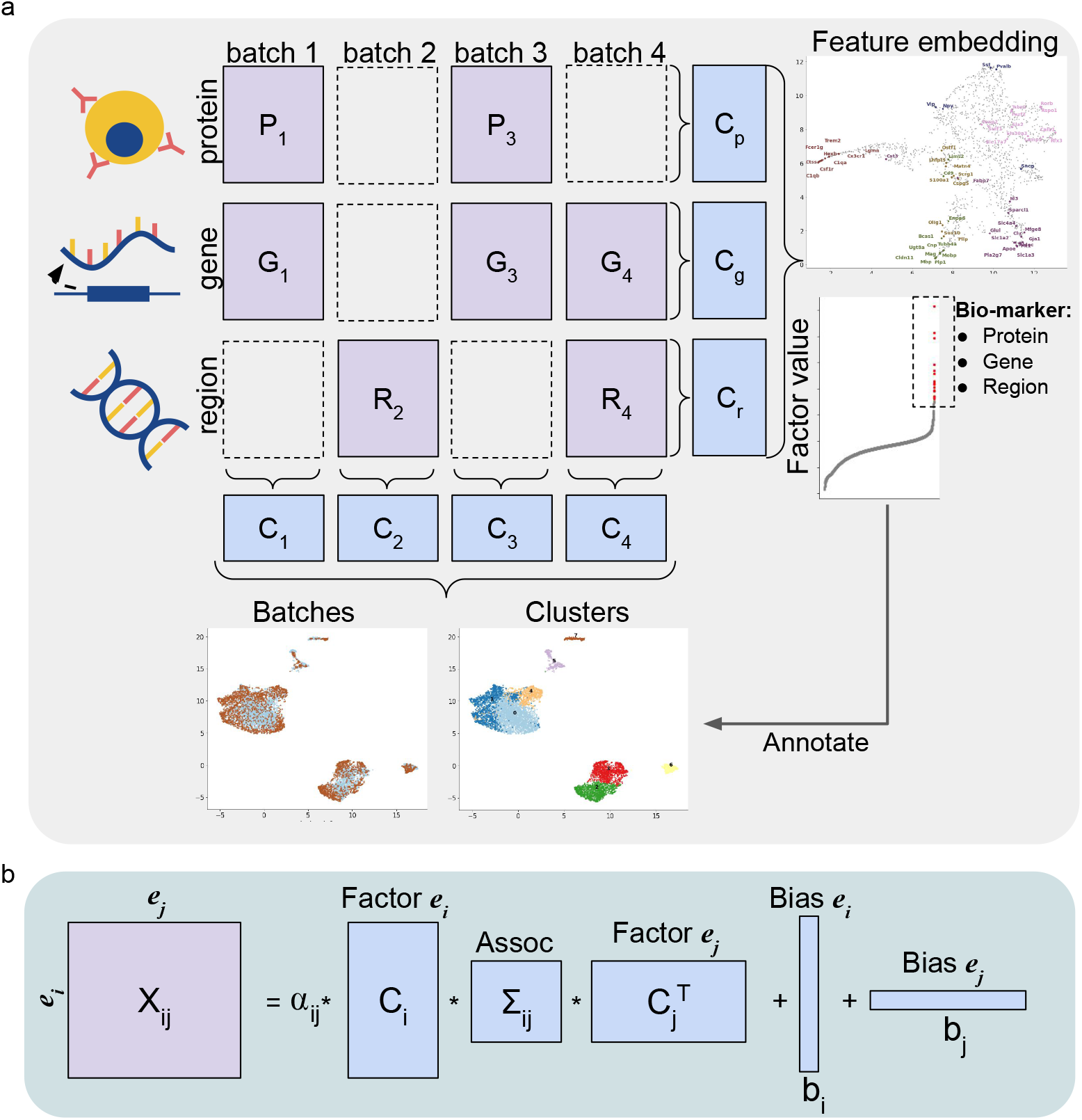
scMoMaT overview. **a**. Graph illustration of running scMoMaT on an example dataset (4 batches and 3 modalities). Given data matrices in a mosaic layout to integrate, scMoMaT outputs cell representations and feature representations of multiple modalities, cell clusters and top-scoring bio-markers of every input modality. The top-scoring markers are used to annotate cell types for the clusters on the learned cell embedding. **b**. Graph illustration showing the factorization of matrix **X**_*i j*_ in scMoMaT.

We tested scMoMaT on both real and simulated datasets covering various kinds of integration tasks. We first test scMoMaT on multiple simulated datasets and quantitatively evaluate its performance. We then test scMoMaT on four real datasets covering drastically different real-life integration tasks, including one human PBMC dataset, one mouse brain cortex dataset, one human bone marrow dataset, and one mouse spleen dataset. We compared the performance of scMoMaT with state-of-the-art data integration methods using multiple benchmarking metrics. The results show that scMoMaT has superior performance in learning cell embedding, and dealing with disproportionate cell type composition between batches. We demonstrated how the multi-modal bio-markers we learned can be used to annotate cell types of the clusters obtained in the integrated space. We also show that these annotations can be better than the annotations provided by the original paper.

## 2 Results

### 2.1 Framework of scMoMaT

scMoMaT uses a matrix tri-factorization framework, which treats each single cell data matrix as a relationship matrix between the “cell” and “feature” entities. A *feature* comes from a modality, which can be gene, region, or protein. Given a single-cell data matrix **X**_*i j*_ from the *i*th cell batch and *j*th feature modality, matrix tri-factorization decomposes it into a cell factor **C**_*i*_, a feature factor **C** _*j*_, and an association matrix Σ_*i j*_. We add bias and scaling terms into the formulation to accommodate the cell- and feature-specific bias and scaling of the data matrix. The objective function for one data matrix is:

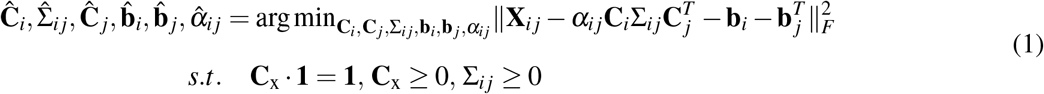

where **b**_*i*_ and **b**_*i*_ are 1 dimensional bias vectors for cell batch *i* and feature modality *j* respectively. *α*_*i j*_ is the matrix scaling parameter. **C**_x_ · **1** = **1, C**_x_ ≥ 0 constrains each row of the factor (**C**_*i*_ and **C** _*j*_) to be a *simplex*, i.e. each row should be non-negative and sum up to 1. This constraint is motivated as follows. In order for the factorization to only capture the major biological variation within the data, the number of latent dimensions *d* (number of columns in **C**_*i*_ and **C** _*j*_) should be much smaller than the number of cells or features in the data matrix. We assume that each latent dimension encodes a distinct biological factor of the dataset, and the factor values of each cell or feature (row vectors of **C**_*i*_ or **C** _*j*_) encode the proportion of each biological factor with the cell or feature identity. In addition, we constrain Σ_*i j*_ to be non-negative. A graphical illustration of the factorization is shown in Fig. 1b.

When integrating multiple data matrices, we construct a loss function with multiple tri-factorization terms (Eq. 1), where each input data matrix corresponds to a tri-factorization term. We use the scenario where the data matrices are from mutiple batches and three modalities: gene, chromatin region and protein as an example (Fig. 1a). Denote the gene expression matrices as 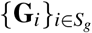, the chromatin accessibility matrices by 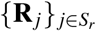, and the protein abundance matrices by 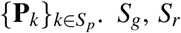, *S*_*g*_, *S*_*r*_ and *S*_*p*_ are the sets of batch indices where gene expression, chromatin accessibility, and protein abundance matrices are available, respectively. The optimization problem of scMoMaT is:

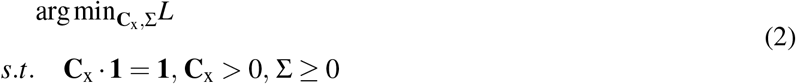

And

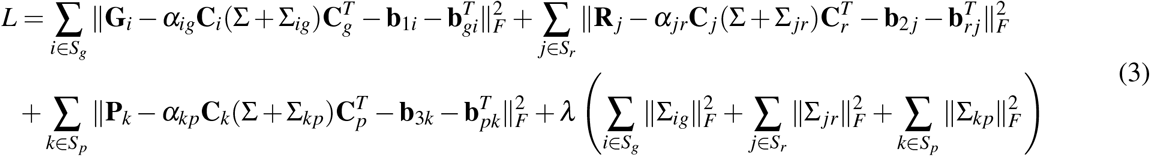

where **C**_*i*_s are the factors for cell batches that have gene expression matrices, **C** _*j*_s are the factors for cell batches that have chromatin accessibility matrices, and **C**_*k*_s are the factors for cell batches that have protein abundance matrices. **C**_*g*_, **C**_*r*_ and **C**_*p*_ are the factors for genes, regions, and proteins. The factors of the same cell batch or feature modality are shared across the tri-factorization terms. Σ is the shared association matrix across all data matrices, and Σ_*ig*_, Σ _*jr*_, Σ_*kp*_ are data matrix-specific association matrices. **b**_xx_s are the cell or feature specific bias vectors for each data matrix. *α*_*ig*_, *α*_*jr*_, *α*_*kp*_ are data matrix-specific scaling parameters. *λ* is the weight that regularize how much data matrix-specific association matrix should vary.

The aforementioned framework works for the cases where there exist common feature modalities across batches (Fig. S1a). When there is no common modality across batches (Fig. S1b), we add pseudo-count matrices (or referred to as *gene activity* matrices in some literature^6^) to make the corresponding modality shared across all batches. We can make the gene modality to be the common modality, and calculate a pseudo-scRNA-seq matrix from scATAC-seq matrix. The procedure to calculate pseudo-scRNA-seq is similar to that used in Seurat v3 and LIGER (Methods).

After the factors are learned by minimizing the objective function, we include an additional post-processing step on the learned cell factors (Methods). The post-processing step constructs a neighborhood graph of all cells, which can be visualized using UMAP and clustered using Leiden cluster algorithm^20^. After obtaining the cluster result of the cells, we retrain the model to learn the feature factors and association matrices. *Feature scoring matrices*, which represent the importance of a feature for a cluster, can then be obtained from the retrained feature factors and association matrices (Methods). These matrices have each latent dimension corresponding to one specific cell cluster, and can be used to extract the cluster-specific top-scoring features (bio-markers) across modalities that jointly define cell type identities.

### 2.2 Testing scMoMaT on simulated datasets

First, we used simulated datasets to test scMoMaT, which allow us to generate different integration scenarios and quantitatively evaluate the integration method. The simulator that we used was similar to the simulator described in scDART^10^, except that continuous cell populations were generated in scDART^10^, whereas clusters of cells were generated in our tests. The simulation procedure can generate paired scRNA-seq and scATAC-seq data (both modalities are profiled within each cell) from any number of batches (Methods).

#### 2.2.1 Data simulation

We generated 6 batches of paired scRNA-seq and scATAC-seq data, which results in 12 data matrices in total. To account for the randomness in the simulation, we repeat the simulation 8 times with 8 different random seeds and report summary results on the 8 datasets. For each dataset, there are 16 cell types shared across batches.

From these 12 data matrices, we created two different integration scenarios, as shown in Figs. 2a,b. In Fig. 2a, no simultaneously profiled cell batch exists. We selected only the scATAC-seq matrices from batches 1, 2, and 3, and selected only scRNA-seq matrices from batches 4, 5, and 6 (totally 6 selected data matrices). In Fig. 2b, there exists one batch of simultaneously profiled cells. We selected scATAC-seq matrices from batches 1, 2, 3 and 4, and scRNA-seq matrices from batches 4, 5 and 6. In order to create unequal cell type compositions across batches, we randomly selected 4 (out of 16) cell types for each data batch and removed these 4 cell types from the batch.

**Figure 2.**
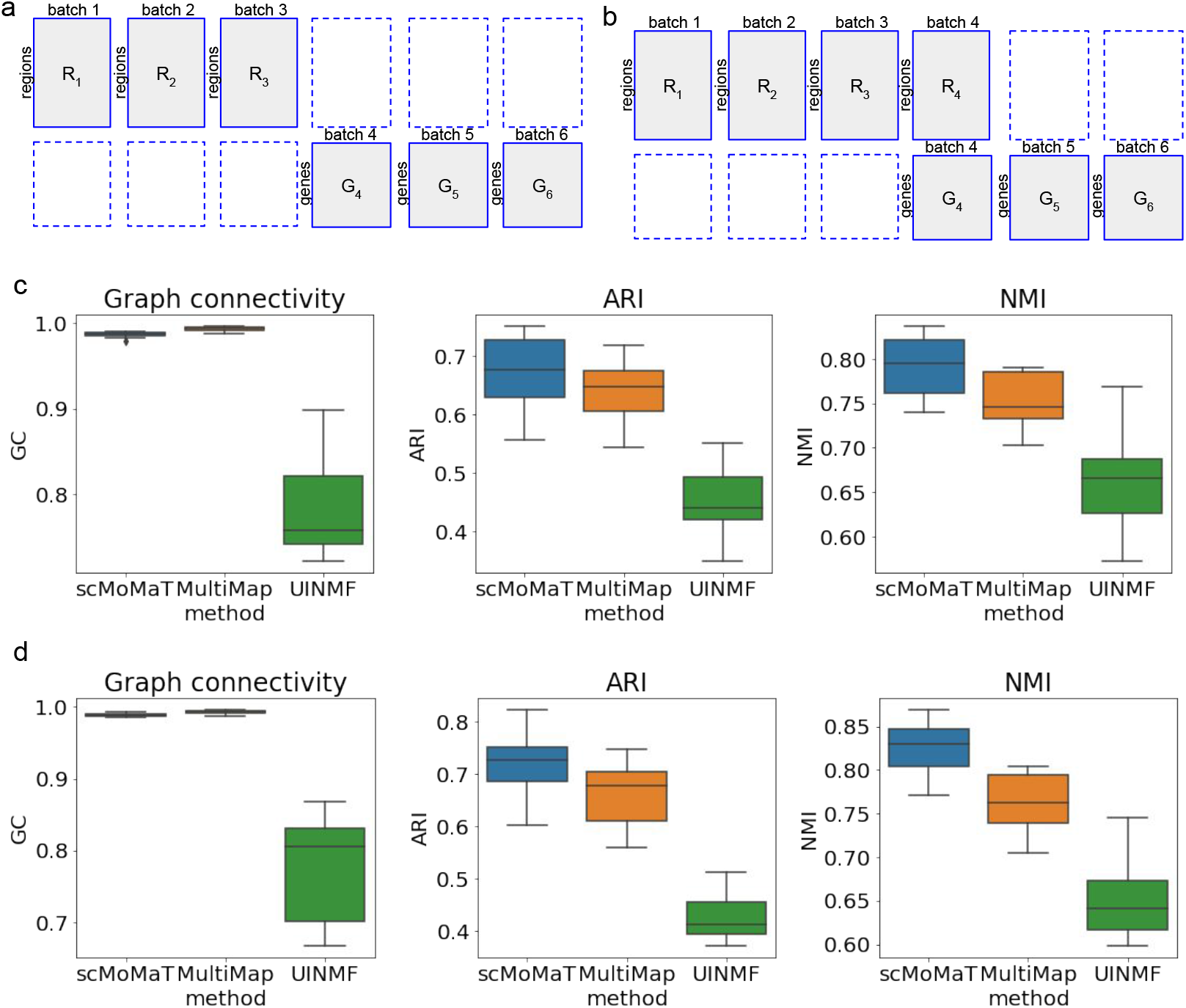
Results on simulated datasets. **a**. The layout of data matrices under the first integration scenario. **b**. The layout of data matrices under the second integration scenario. **c**. Graph connectivity, ARI, and NMI scores of scMoMaT and baseline methods under the first integration scenario. **d**. Graph connectivity, ARI, and NMI scores of scMoMaT and baseline methods under the second integration scenario.

#### 2.2.2 Results on simulated data

We compared the performance of scMoMaT with two recently published methods which can work with these integration scenarios: MultiMap^21^ and UINMF^16^. We ran scMoMaT, UINMF and MultiMap under the first integration scenario. Firstly, we filled in the missing scRNA-seq matrices of the first three batches with pseudo-count matrices (Methods). Then, we ran scMoMaT and set its latent dimension *d* = 20, number of neighbors *k* = 30, and radius parameter *r* = 0.7 for all runs. Details and parameter settings of MultiMap and UINMF are included in Methods.

We quantitatively measured the overall performance of three methods with three scores: k-nearest neighbor graph connectivity (kNN-GC or GC), normalized mutual information (NMI), and adjusted Rand index (ARI) (Methods). These metrics were used in^22^ to benchmark various integration methods, where GC measures batch effect removal per cell identity label, and NMI and ARI measure conservation of biological variation during integration in terms of cell identity labels.

We summarized the performance of each method on 8 datasets using boxplots (Fig. 2c). The results show that scMoMaT performs comparably with MultiMap in matching cell batches (similar GC score), and perform consistently better in separating cell types (higher ARI and NMI scores). We visualized the latent embedding of scMoMaT and baseline methods on one of the 8 datasets using UMAP (Fig. S2), and the visualization shows that with scMoMaT the cell types are better separated, and the locations of the same cell type in the UMAP space are more consistent across batches. Taking cluster 16 which is missing in batches 2 and 3 as an example: in the results of MultiMap, cluster 16 is at consistent locations in batches 1, 4, 5, 6, but in batches 2 and 3, some cells from other clusters are placed at this location (circled in red). In the results of UINMF, cluster 16 in batch 1 is located in a different area from that in batches 4, 5, 6 (circled in red).

We then measured the performance of all three methods under the second integration scenario (as shown in Fig. 2b). We filled in the missing scRNA-seq matrices for the first three batches with pseudo-count matrices (the same as the first test scenario), and ran methods with hyper-parameter settings the same as the first test scenario. To make MultiMap applicable to the dataset, we concatenated the scATAC-seq and scRNA-seq in cell batch 4 into a single data matrix and reduced the dimensionality of that batch by running PCA on the concatenated matrix.

Boxplots of GC, ARI and NMI scores of each method on 8 datasets are shown in Fig. 2d. The boxplots again show a better overall performance of scMoMaT compared to the two baseline methods.

### 2.3 scMoMaT performs mosaic integration on human PBMC dataset and annotates sub-cell-types

We applied scMoMaT to a human PBMC dataset which includes 4 batches of cells^3^. The first 2 batches of cells are measured with gene expression and protein abundance simultaneously using CITE-seq^23,24^ (batch 1 has 5023 cells, and batch 2 has 3666 cells); The last 2 batches of cells are measured with protein abundance and chromatin accessibility simultaneously using ASAP-seq^3^ (batch 3 includes 3517 cells, and batch 4 includes 4849 cells). In total, there are 8 data matrices (Fig. 3a). On this dataset, we demonstrate: (1) scMoMaT produces integration with higher quality in terms of preserving cell identity and mixing batches compared to baseline methods; (2) scMoMaT improves cell type annotation through integration; (3) The bio-markers learned from multiple modalities lead to cell type annotation with higher resolution.

**Figure 3.**
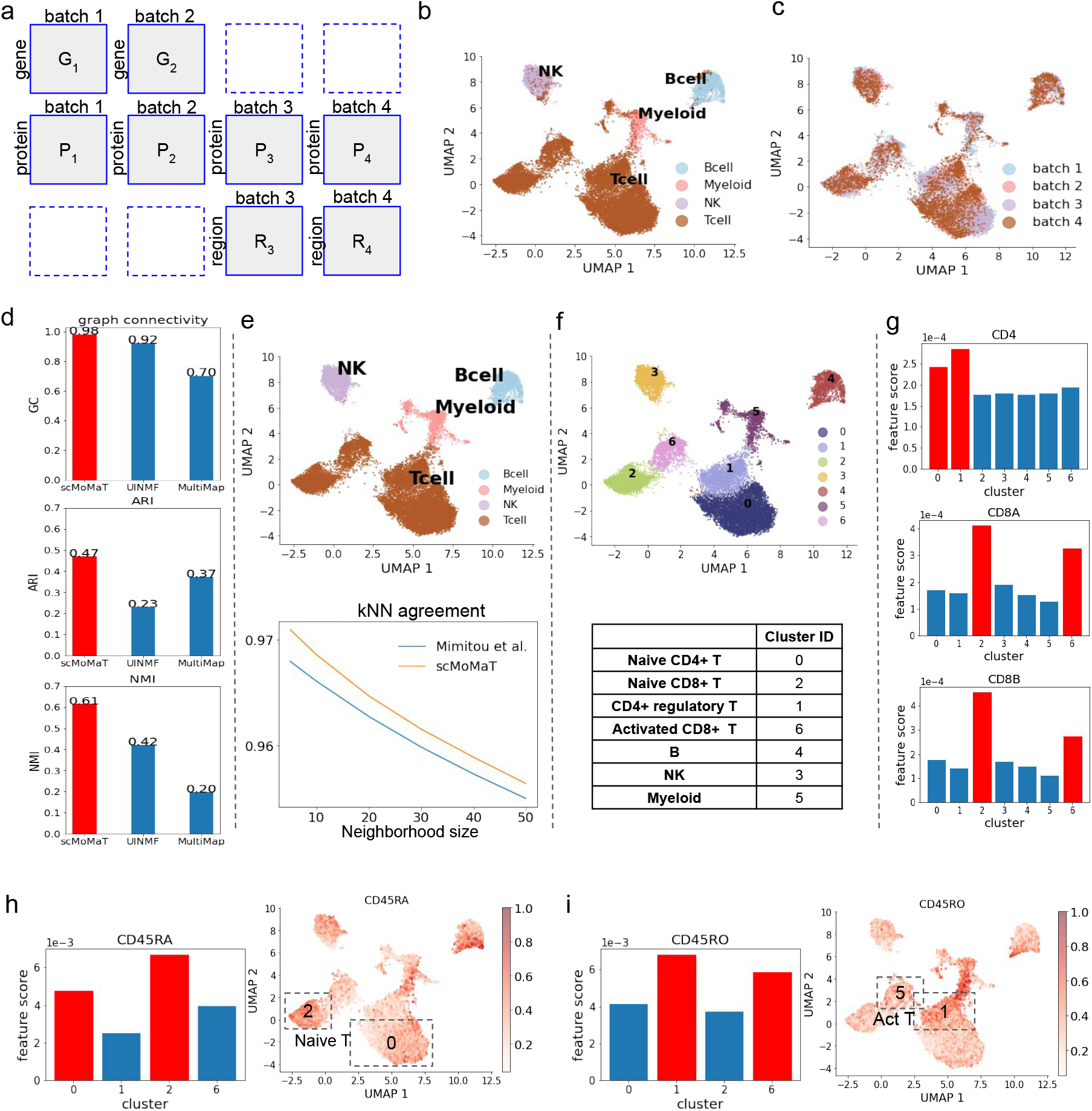
Results on the human PBMC dataset. **a**. Layout of input data matrices in human PBMC dataset. **b**-**c**. The UMAP visualization of cell factors learned by scMoMaT, where cells are colored by (**b**) cell type labels from the original data paper (Mimito et al) and (**c**) data batches. **d**. The graph connectivity, ARI, and NMI scores of scMoMaT and baseline methods. The top-scoring method is colored red. **e**. (Upper) The UMAP visualization of cell factors, where cells are colored by the label from scMoMaT. (Lower) The kNN agreement scores of scMoMaT labels and labels in original data paper (Mimito et al) under different neighborhood sizes *k*. **f**. (Upper) The UMAP visualization of cell factors; cells colored by Leiden clustering labels. (Lower) Cell type annotation for the Leiden clusters. **g**. scores of marker genes *CD4, CD8A*, and *CD8B* in different clusters, where x-axis correspond to Leiden clusters. The top-scoring clusters are colored red. **h**,**i**. The barplots show the scores of marker protein *CD45RA* and *CD45RO* in different clusters. The heatmaps show the abundance level of proteins *CD45RA* and *CD45RO*, where the top-scoring clusters are annotated in frames.

First, we visualized the cell factors of all batches learned by scMoMaT using UMAP (Figs. 3b,c). In Fig. 3b, the cells are colored with the cell type labels obtained from the original data paper^3^, where different cell types are overall separated. In Fig. 3c, cells are colored by batches and cells from different batches are mixed in the integrated data. We compared the performance of scMoMaT with MultiMap^21^ and UINMF^16^ (Details and parameter settings are included in Methods).

We measured the performance using metrics including GC, ARI and NMI scores. Cell type labels provided in the original data paper^3^ were used as ground truth clustering labels. The results (Fig. 3d) show that scMoMaT performs better than the two baseline methods with all metrics. Indeed, visualizations of latent embedding from MultiMap and UINMF (Figs. S3a,b) show that the cell types were not matched correctly between different batches for both methods.

Although it is a standard practice to test how much integration methods preserve the cell identity annotated in the original paper of the dataset (as shown in Fig. 3d)^21,22^, we take a step further and ask whether we can improve this cell identity annotation through integrating data from multiple modalities. Performing Leiden clustering on the cell representations learned by scMoMaT, we obtained the clusters shown in Fig. 3e (upper plot) and mapped the labels to the clusters. We set to compare the labels from the original paper by Mimitou *et al* and those obtained from scMoMaT. We consider that good cell labels should be consistent with the cell-cell variation in every batch and every modality. In Fig. S4 we visualize each input data matrix respectively with the cell labels from scMoMaT and Mimitou *et al*. Visually, although in most of the plots different cell types are separated in the UMAP space, there are areas where more than one cell types are mixed (eg. the areas circled in red). To quantify which set of labels has better agreement with the cell-cell variation in individual data matrix, we used a metric named *k-nearest neighbor agreement* (kNN agreement). For each cell, this metric measures the percentage of cells that have the same label as the given cell in its *k* nearest neighbors (Methods). Fig. 3e (lower plot) shows the kNN agreement score of each set of labels averaged over all cells in all input data matrices, where the scMoMaT labels have improved over the original labels from Mimitou *et al*.

We then show that the feature factors learned by scMoMaT give rise to bio-markers from multiple modalities which can be used to annotate cell types at higher resolution. We ran Leiden clustering algorithm on the integrated latent space of cells and obtained seven clusters (Fig. 3f, upper plot). We then fed the cluster labels into scMoMaT for retraining to obtain feature scoring matrices, which show the importance score of features in each cluster. Therefore, for each cell cluster, we have three vectors: (1) a vector representing the importance of each gene for this cluster; (2) a vector representing the importance of each chromatin region for this cluster; and (3) a vector representing the importance of each protein for this cluster. After we included the motif deviation matrix from chromVAR^17^ analysis, we were also able to obtain vectors representing the importance of each motif for every cluster. The top scoring features (genes, chromatin regions, proteins, and motifs) in these vectors can be used as bio-markers for cell type annotation.

The complete annotation of these clusters is shown in Fig. 3f (lower table). We now discuss how the top scoring features (with the highest importance score) from each modality lead to this annotation. First, we annotate clusters 3, 4 and 5 using top-scoring genes. The top-20 genes for cluster 3 include *GNLY, NKG7, KLRD1*, and *KLRF1*, which are the marker genes of Natural Killer (NK) cell^25,26^. The top-20 genes in cluster 4 include *MS4A1, CD79A* and *CD37*, which are the marker genes of B cells^26,27^. The top-20 genes of cluster 6 includes *CTSS*^28^, *SPI1*^29^, *and CD63*^30^, which are the marker genes of Myeloid cells (Fig. S5a, marker genes with red frames). These annotations are further confirmed by extra marker genes for these cell types from CellMarker^31^ (Fig. S5a, known marker genes in blue frames). Furthermore, these annotations are consistent with the annotations from the original paper in visualization (Fig. 3b). These evidence together show that the top-scoring genes learned by scMoMaT are highly consistent with known knowledge.

We have higher scores of CD3G, CD3E and CD3D (which are T cell markers) in clusters 0, 1, 2, 6 than other clusters (Fig. S5b), so we tentatively annotate these clusters as T cells. This is in agreement with the annotation in Fig. 3e. The feature factors learned by scMoMaT can be used to further identify T cell subtypes in the integrated data. First, clusters 0, 1 have higher *CD4* scores, which shows that they correspond to CD4^+^ T cells. Clusters 2, 6 have higher *CD8A* and *CD8B* scores, which shows that they correspond to CD8^+^ T cells (Fig. 3g). The distributions of the expression value of *CD4, CD8A*, and *CD8B* also matches the importance scores of these genes (Fig. S6a).

Within CD4^+^ and CD8^+^ T cells, top scoring proteins can be used to further separate them into naive T cells and activated T cells. Naive T cells have high abundance of surface protein *CD45RA* and low abundance of surface protein *CD45RO*. Activated T cells, on the contrary, have high *CD45RO* and low *CD45RA*^32,33^. *Using the scores of these proteins, we annotate clusters 0 and 2 to be naive T cells (lower CD45RO* score and higher *CD45RA* score, Fig. 3h), and cluster 1, 6 to be activated T cell (higher *CD45RO* score and lower *CD45RA* score, Fig. 3i). The high importance scores of Naive T cell marker genes (*CD27, TCF7*) in clusters 0 and 2 also confirms the annotation of Naive T cell from protein scores^33,34^ (Fig. S6b). Cluster 1 is further shown to correspond to CD4^+^ regulatory T cell (Treg) using the importance scores of marker genes *Foxp3*, and *IL2RA*^34^ *(Fig. S6c). Cluster 6 has high scores of cytotoxicity markers GZMK* and *GZMB*^34^ (Fig. S6d), which further confirms the activated CD8^+^ identity. Using the marker gene information in CellMarker^31^, we found extra marker genes for the cell type annotated to clusters 0, 1, 2, 6 from the top-20 genes of these clusters (Fig. S6e, with known marker genes in blue frames).

These discussions all together lead to the final annotations shown in Fig. 3f. The annotations are further confirmed by the known protein markers in the top scoring proteins (Fig. S7, with marker proteins in blue frames). Because scMoMaT also incorporates the motif deviation matrix learned by chromVAR^17^ from the scATAC-seq data matrix, scMoMaT also outputs top-scoring motifs for each cluster. The known motif markers in the top-scoring motifs also confirm our cell type annotations (Fig. S8, source of motif markers in Table S1).

### 2.4 scMoMaT mosaic integration on mouse cortex data

We then applied scMoMaT on a mouse brain cortex dataset. We collected 5 batches of mouse brain cortex datasets from different publications. The first data batch has 10, 309 cells where chromatin accessibility and gene expression were simultaneously measured using SNARE-Seq^2^. The second batch measures the gene expression of 40, 166 cells using 10x v3 single-nucleus RNA-Sequencing technology (snRNA-seq) and the third batch measures the chromatin accessibility of 8718 cells using single-nucleus ATAC-Sequencing (snATAC-seq)^35^. The fourth batch measures the gene expression of 14249 cells and is obtained from Allen Brain Atlas^36,37^. The fifth batch measures the chromatin accessibility of 3512 cells and is obtained from 10x Genomics website. In total, 6 data matrices are used as input to scMoMaT and they are organized as Fig. 4a.

**Figure 4.**
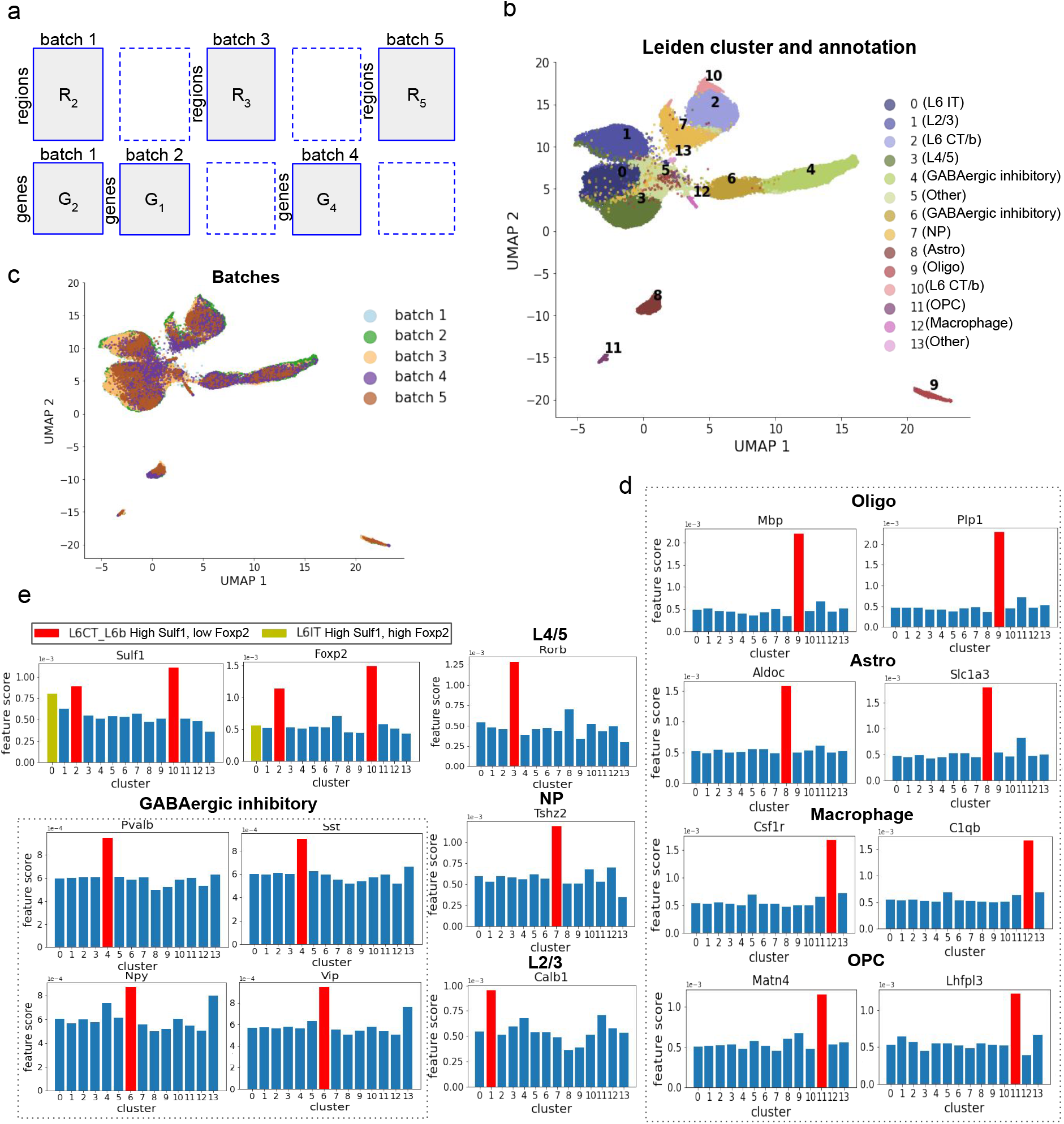
Results on mouse brain cortex dataset. **a**. Layout of input data matrices in mouse brain cortex dataset. **b**,**c**. The UMAP visualization of cell factors learned by scMoMaT, where cells are colored by (**b**) Leiden clusters (with scMoMaT-annotated cell types) and (**c**) cell batches. **d**. The scores of marker genes for non-neuronal cell types, where x-axis correspond to Leiden clusters. **e**. The scores of neuronal cell type marker genes in different clusters. The top-scoring clusters are colored red.

First, to understand the variation structure between cells in each data matrix before integration, we visualized each data matrix separately using UMAP. The cells are colored using the cell type labels curated and re-organized from the original data paper (Methods, Fig. S9). The visualizations show differences in the variation structures between different batches and modalities, which can be caused by various factors, including technical confounders such as read depth and noise level^38,39^, or disparity of cell type composition between batches. Applying scMoMaT to these matrices can lead to integrated cell representations in a shared latent space, and the top scoring features output from scMoMaT can be used as bio-markers for cell type annotation. Through the latent space representations of genes and regions learned by scMoMaT, we also demonstrate that bio-markers for the same cell type tend to have similar low-dimensional representations.

scMoMaT took as input the six data matrices and two additional pseudo-scRNA-seq matrices that were calculated from scATAC-seq matrices for data batches 3 and 5 (Methods). The learned cell representations are shown in Figs. 4b-c, where the clusters in Fig. 4b were obtained by running Leiden clustering^40^ on the cell factors, and the cell types were annotated with bio-markers learned by scMoMaT. The cell type annotation process is described below.

We first collected known marker genes for the cell types included in these datasets from existing literature^35,41,42^ (Table S2). The scores of these marker genes learned through the retraining step were used to annotate the clusters in Fig. 4b: for the non-neuron cell types, *Mbp* and *Plp1* annotate cluster 9 as oligodendrocyte (Oligo), *Aldoc* and *Slc1a3* annotate cluster 8 as Astrocytes (Astro), *Csf1r* and *C1qb* annotate cluster 12 as Macrophage, *Matn4* and *Lhfpl3* annotate cluster 11 as oligodendrocytes (OPC) (Fig. 4d). For the neuronal cell types, L6 neuron marker gene *Sulf1* has high scores in clusters 0, 2, and 10 (Fig. 4e). Out of these three clusters, *Foxp2* was used to distinguish L6 corticothalamic neuron (L6 CT/b, cluster 2 and 10, with high *Foxp2*) from L6 intratelencephalic neuron^35^ (L6 IT, cluster 0, with low *Foxp2*, Fig. 4e). The scores of *Rorb* annotate cluster 3 as L4/5 excitatory neuron^35^, *Tshz2* annotate cluster 7 as near-projecting excitatory neurons (NP), and *Calb1* annotate cluster 1 as L2/3 excitatory neurons (Fig. 4e). We also found high scores of marker genes *Pvalb, Sst, Npy, Vip* in clusters 4 and 6, which shows that those two clusters corresponds to GABAergic inhibitory neurons (Fig. 4e). Cluster 4 has higher scores of *Pvalb* and *Sst* and cluster 6 has higher scores of *Npy* and *Vip*, which shows that these two clusters correspond to distinct sub-cell types in GABAergic inhibitory neurons^43^ (Fig. 4e).

The top-20 scoring genes for each cluster are enriched with known marker genes for the annotated cell type, according to marker genes collected in Table S2 and in CellMarker^31^ (Fig. S10, with known marker genes in blue frames). There are fewer marker genes found for neuronal cell subtypes partly because fewer marker genes are known for these cell types.

Including the motif deviation matrix (from chromVAR) allows scMoMaT to learn top scoring motifs for each clusters (Fig. 5a). Out of the top-20 motifs, we see MA0062.2_Gabpa, MA0117.2_Mafb and MA0002.2_RUNX1 for Macrophage (cluster 12), MA0515.1_Sox6, MA0442.1_SOX10 and MA0514.1_Sox3 for oligodendrocyte (cluster 9), MA0463.1_Bcl6 and MA0518.1_Stat4 for L6 CT/b neuron (clusters 2 and 10), etc^44^. In particular, motifs MA0623.1_Neurog1 and MA0461.2_Atoh1 stand out in L6 CT/b, and *Neurog1* and *Atoh1* are reported to be important transcription factors in neurogenesis^45,46^. Overall, these motifs further support our cell type annotations.

Since scMoMaT jointly learns the region and gene factors along with the cell factors, we also visualize the region and gene factors (Figs. 5b,c). Fig. 5b shows the gene factors where known marker genes for different cell types are marked with different colors. One can observe that the marker genes for GABAergic inhibitory neurons, oligodendrocyte, oligodendrocyte precursors, Macrophage, Astrocytes, and Glutamatergic neurons (including L2/3, L4/5, L6 IT, L6 CT/b, NP) are clearly separated into different areas of the UMAP space. For the region factors, we map a region to a gene if the region is located within the 2000 base-pair upstream or the gene body of the gene on the genome, and we plot the genes as the average of the regions associated with the corresponding gene (genes are represented by colored dots in Fig. 5c). Fig. 5c shows that the chromatin regions that correspond to marker genes of oligodendrocyte & oligodendrocyte precursors, Macrophage, and Glutamatergic neurons are also separated into distinct areas based on the region factors. Both the gene and the region factors show that genes and regions do not form distinct clusters (which is expected because genes like house keeping genes do not belong to a specific gene module), but marker genes of different cell types are separated in the gene and region factor space learned by scMoMaT.

**Figure 5.**
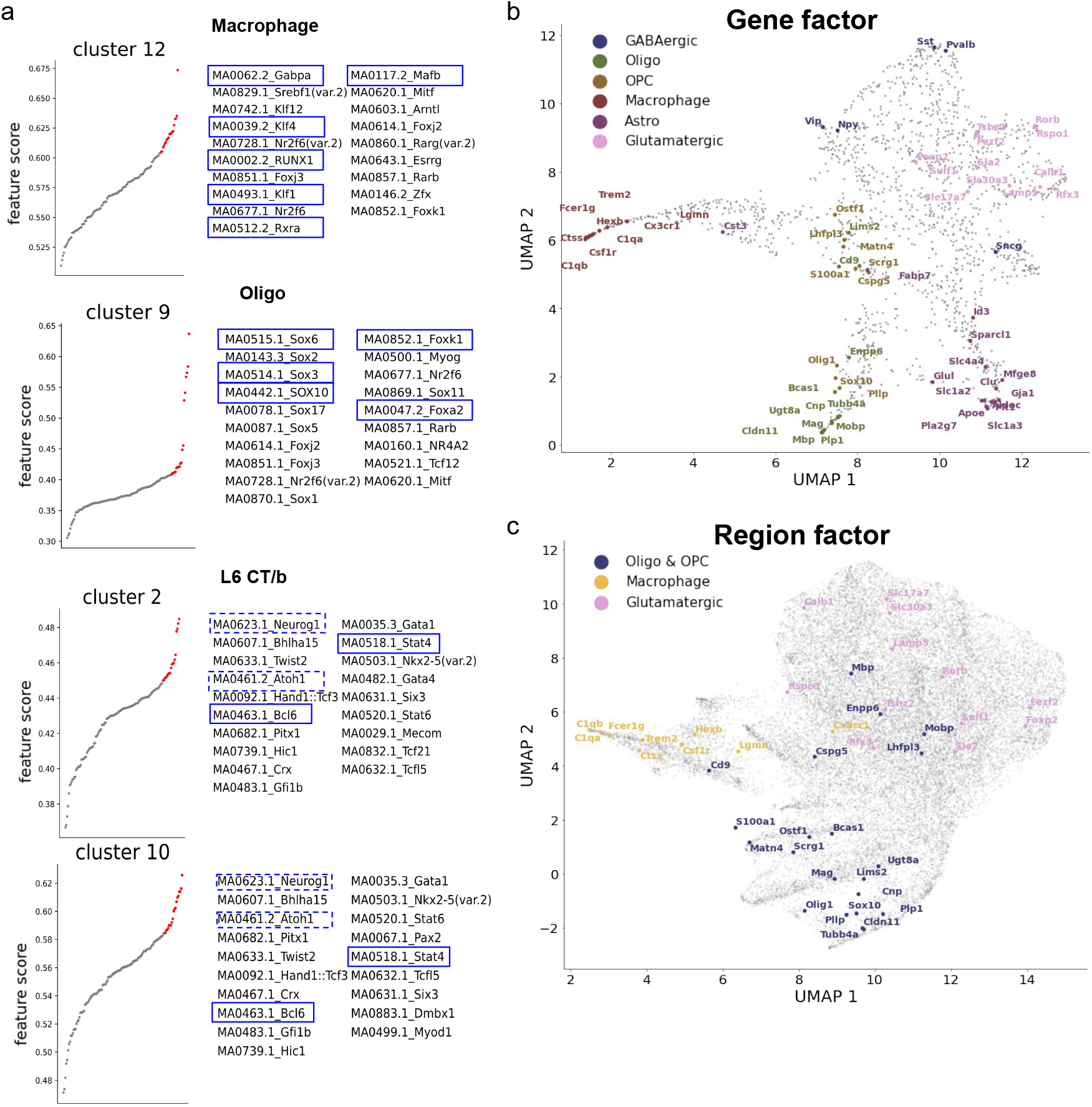
Additional results on mouse brain cortex dataset. **a**. The top-20 scoring motifs in cluster 12 (Macrophage), 9 (Oligo), and 2 (L6 CT/b). Motifs with TFs reported for a specific cell type are highlighted in blue frames. **b**,**c**. The UMAP visualization of (**b**) gene factors and (**c**) region factors; known marker genes of different cell types are annotated with corresponding colors.

### 2.5 scMoMaT integrates batches with no shared modalities

It is a very challenging integration scenario if the batches do not share any modality (also called diagonal integration). The most common example of such a scenario is the integration of a scATAC-seq matrix and a scRNA-seq matrix obtained from different batches. To integrate such datasets, additional assumptions or information often need to be provided. Some methods assume that the latent distributions of cells are similar between batches, which fails to accommodate the cases where the data batches have unequal cell type compositions. Other methods transform the scATAC-seq matrix into a pseudo-scRNA-seq matrix using the cross-modalities relationship, and integrate the scRNA-seq matrix and pseudo-scRNA-seq matrix. Using the pseudo-scRNA-seq instead of the scATAC-seq matrix, these methods may suffer from the errors introduced during the process of calculating the pseudo-scRNA-seq matrix and do not fully utilize the epigenomic information in the scATAC-seq matrix. scMoMaT, on the other hand, keeps both the original scATAC-seq matrix and the pseudo-scRNA-seq matrix in order to better exploit the scATAC-seq information. Also, we binarized the pseudo-scRNA-seq matrices as a denoising step (Methods).

We applied scMoMaT to a healthy human bone marrow mononuclear cells (BMMC) dataset^47^. The dataset includes two batches of cells, where the first batch has 16510 cells sequenced with scATAC-seq and the second batch has 12601 cells sequenced with scRNA-seq (matrix relationships follows Fig. 6a). scMoMaT takes as input both matrices, and generates a pseudo-scRNA-seq matrix for the second batch using its scATAC-seq data matrix (Methods). We visualize the cell factors learned from scMoMaT (Figs. 6b,c) using UMAP, and color the cells using the literature-derived labels (Fig. 6b) and data batches (Fig. 6c). In the visualizations, cell batches are well integrated in the latent space, while cell identities in each data batch are also preserved.

**Figure 6.**
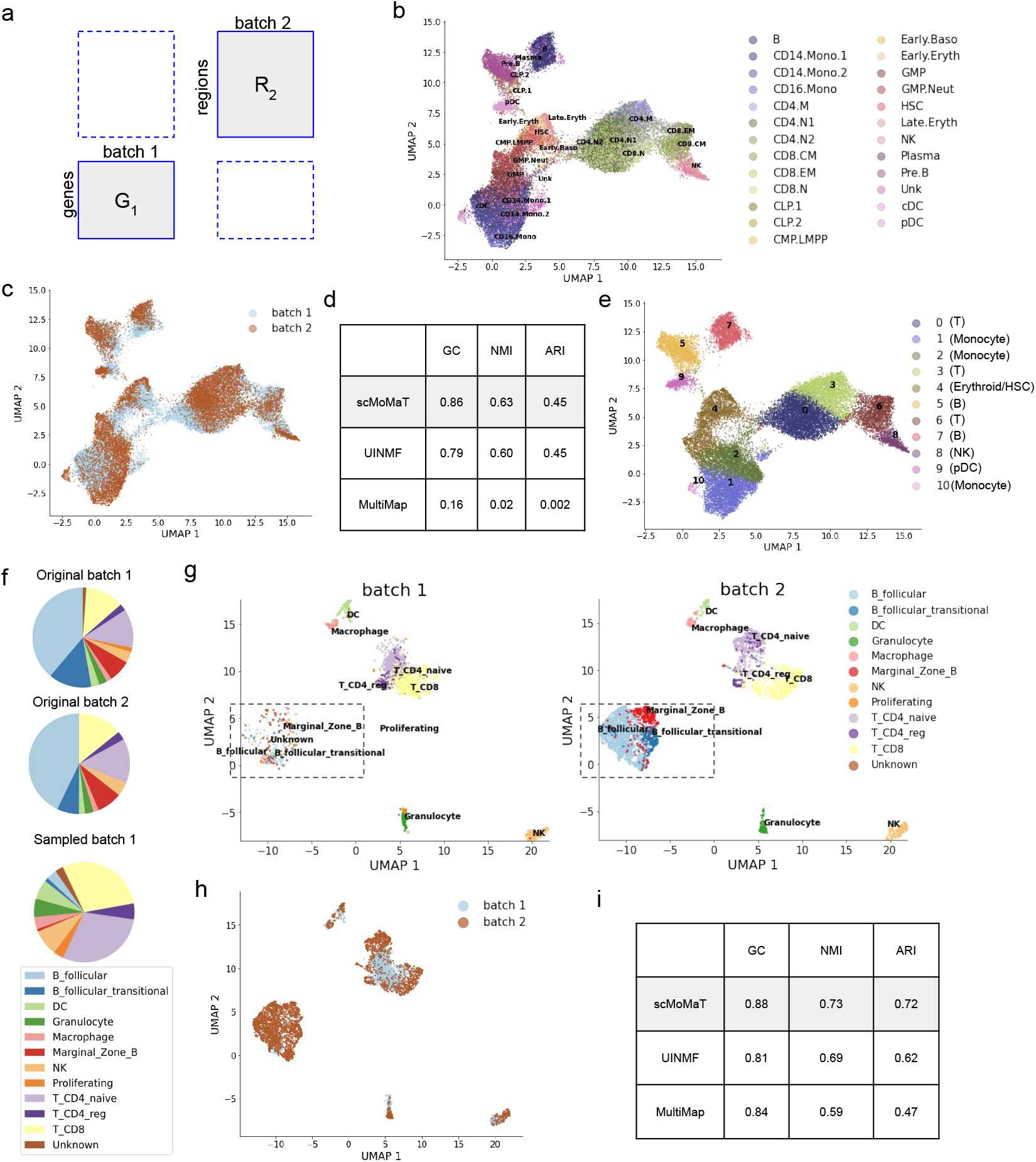
Results on the human bone marrow and mouse spleen dataset. **a**. Relationship between data matrices in the two datasets. **b**,**c**. The UMAP visualization of cell factors, where cells are colored by (**b**) ground truth cell type, and (**c**) data batches. **e**. The graph connectivity (GC), ARI, and NMI scores of scMoMaT and baseline methods. **e**. The UMAP visualization of cell factors, where cells are colored by Leiden clusters (with scMoMaT-annotated cell types). **f**. The cell type composition of each batch in original and sub-sampled mouse spleen dataset. **g**. The UMAP visualization of cell factors learned from sub-sampled dataset; Batches 1 and 2 are visualized separately. The disproportionate B follicular cells are annotated in frames. **h**. The UMAP visualization of cell factors learned from sub-sampled dataset, where cells are colored by data batches. **i**. The graph connectivity (GC), ARI, and NMI scores of scMoMaT and baseline methods on the sub-sampled dataset.

We also ran UINMF and MultiMap on the dataset (results visualized in Figs. S11). We quantitatively measured the overall performance of the methods using GC, NMI, and ARI scores (Methods, Fig. 6d). scMoMaT has the highest GC score, which shows that scMoMaT better matches the same cell type between batches. scMoMaT and UINMF have similar ARI and NMI scores. MultiMap, on the other hand, mixes cells from different cell types in the latent space. It may be due to the fact that the cell types in BMMC dataset are closely located in the original dataset as they follow the trajectories of the hematopoiesis process, and MultiMap fails to distinguish the closely located cell types (Fig. S11b).

After clustering the cells, scMoMaT learned the importance scores of genes and regions in each cluster. We annotated cell types according to the top scoring genes and regions. The cluster result and cell type annotations are shown in Fig. 6e. Since we also input the motif deviation matrix obtained by chromVAR, scMoMaT learns top scoring motifs along with genes and regions. Multiple known marker genes and relevant motifs are shown to have high scores for their corresponding cell types. In cluster 8 (natural killer (NK) cells), scMoMaT found a high score of marker gene *GNLY*^25,26^, which matches the gene expression pattern in the dataset (Fig. S12a). Similarly, scMoMaT found T-cell marker gene *CD3D*^34^ in clusters 0, 3, and 6 (Fig. S12a), B-cell marker gene CD79A^27^ in clusters 5 and 7 (Fig. S12a), Monocyte marker gene *S100A9*^48^ in clusters 1, 2, and 10 (Fig. S12a), and Plasmacytoid Dendritic Cell (pDC) marker gene *PTPRS*^49^ in cluster 9 Fig. S12a). The top-20 genes of clusters 0, 3, and 6 reveal even more T-cell-related marker genes including *BCL11B, IL7R, LEF1*, etc^31^ (top-20 genes in Fig. S12b, with known marker genes in blue frames).

Meanwhile, the top motifs for each cluster also confirmed the cell type annotations. scMoMaT found high scores of motif *MA0102*.*3_CEBPA, MA0837*.*1_CEBPE*, and *MA0466*.*2_CEBPB* in Monocytes. Their corresponding tran-scriptin factors *CEBPA, CEBPE*, and *CEBPB* are known to be monocyte-differentiation regulators^47,50,51^ (Fig. S13a). In addition, motifs *MA0800*.*1_EOMES, MA0802*.*1_TBR1*, and *MA0690*.*1_TBX21* have high scores in cluster 8 (NK). *EOMES* regulates the maturation of NK cells^52^, whereas *TBX21* (also known as T-bet, belonging to T-box subfamily *TBR1*) is also known to orchestrate the development and effector functions in NK cells^53^(Fig. S13b).

Finally, the cell type annotations obtained with the learned bio-markers are overall consistent with the original cell type annotations (Figs. 6b,e), which verifies that the bio-markers we learned are meaningful.

### 2.6 scMoMaT integrates batches with unequal cell type compositions

In this section, we test how well scMoMaT performs when the cell type compositions are unequal between batches. We used a mouse spleen dataset^21,54^ that includes two batches of cells, where the first batch has 4382 cells sequenced with scRNA-seq, and the second batch has 3166 cells sequenced with scATAC-seq (matrix relationships follows Fig. 6a). The dataset mainly consists of T cells (1190 cells in Batch 1, 990 cells in Batch 2), B cells (2621 cells in Batch 1, 1835 cells in Batch 2), and some other cell types that reside in mouse spleen. The original two data batches have similar cell type compositions (Fig. 6f). We created data batches with disproportionate cell types by sub-sampling the most populated cell type, B cells (including B_follicular, B_follicular_transitional and Marginal_zone_B), in Batch 1 such that only 100 B cells were left. The sub-sampling step changed the proportion of B cells from 59.8% to 5.4%, which drastically changed the cell type composition of Batch 1 (Fig. 6f).

We applied scMoMaT, UINMF and MultiMap to this dataset. The visualization shows that scMoMaT can correctly match cell types in two data batches regardless of the disproportionate cell type compositions between two batches, especially B cells which barely exist in the first batch (Figs. 6g,h). The cell factors of two batches are separately plotted in Figs. 6g for the two batches, where B cells lie within the box. UINMF and MultiMap also perform reasonably well in terms of integrating the two batches (Fig. S14), but the cell types are not clearly separated in MultiMap (Fig. S14a-b, Fig. 6i). All three methods show robust performance towards disproportionate cell type compositions with scMoMaT having the best overall performance among these methods (Fig. 6i).

## 3 Discussion

In this study, we introduced scMoMaT, a single cell data integration method that works on mosaic integration scenario. We applied scMoMaT on different mosaic integration tasks. The results validated the broad applicability of scMoMaT under various types of data integration scenarios. We showed that scMoMaT not only has superior performance compared to existing methods in terms of metrics used to evaluate integration methods, but also learns cluster-specific bio-markers from every input modality that can be used to annotate cell types in the integrated cell space with high confidence. The new annotations can improve the annotations provided in the original papers that publish the datasets, as the clustering and annotations from scMoMaT comprehensively consider information from multiple modalities. Furthermore, we also showed that scMoMaT is able to integrate batches that have disproportionate cell type compositions. With the increasing availability of single cell multi-omics datasets, we expect that scMoMaT will be widely applied to various data integration tasks.

Compared to data integration methods that only learn cell representations in the integrated space, scMoMaT also learns feature representations (eg, gene representations). In the future, considering feature representations in the data integration framework can help with learning cross-modality relationships from single cell multi-omics data.

## 4 Methods

### 4.1 Training procedure of scMoMaT

We minimize the loss function of scMoMaT (Eq. 2 and 3) using mini-batch stochastic gradient descent. Within each iteration, we pick one parameter matrix from cell and feature factors (the **C**_x_ matrices), shared and data matrix-specific association matrices ({Σ, Σ_xx_}), bias matrices (**b**_xx_), and scaling parameter *α*_xx_ and fix the other parameter matrices. Then, we update a mini-batch of the selected parameter matrix using gradient descent. Each mini-batch is constructed by sub-sampling 10% of cells and features in each data matrix. Then we loop through all parameter matrices and update them using gradient descent in order.

In order to enforce the simplex constraint on the factor matrices, we transform the original factor matrices using a softmax function before using it to calculate the reconstruction loss and use the softmax-transformed factor matrices as the output factor matrices of the model. We enforce the non-negativity constraint on the shared association matrix Σ by changing all its negative values to zero every time that it is updated.

In each iteration, we update the bias terms and scaling parameters using closed-form solutions by setting its gradient to 0. Taking the data matrix **G**_*i*_ as an example, the closed-form solution of its cell bias term **b**_1*i*_ follows:

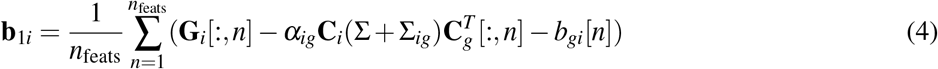

where *n*_feats_ is the total number features in **G**_*i*_. Similarly, its feature bias term **b**_*gi*_ follows:

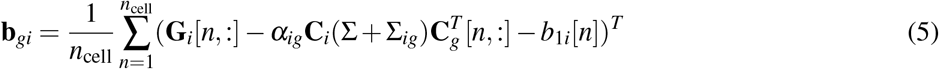

where *n*_cell_ is the total number of cells in **G**_*i*_. The scaling parameter can be calculated as

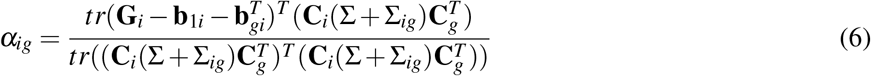

### 4.2 Calculating pseudo-scRNA-seq matrix from scATAC-seq matrix

When the data matrices do not share a common cell batch or feature entity (e.g. diagnol integration scenario), scMoMaT uses the relationship between different feature entities to create pseudo-count matrices that can fill in the positions of missing modalities or batches during integration. When integrating scATAC-seq matrix and scRNA-seq matrix from different batches, scMoMaT creates a pseudo-scRNA-seq for each batch that has only a scATAC-seq matrix. scMoMaT constructs pseudo-scRNA-seq matrix from scATAC-seq matrix by summing up the region counts from all regions that lie within the 2000 base-pair upstream from the TSS of the gene and the regions that lie within the gene body on the genome. Different from the gene activity matrix that was used in Seurat v3 and LIGER, scMoMaT further binarizes the pseudo-scRNA-seq instead of directly using it for integration. The use of additional binarization step is based on two reasons: (1) the relationship between region counts and gene counts is not linear. The activation of the promoting regions of a gene correlate with the activation of the transcription process of the gene, but there is not enough evidence showing that the gene expression level is positively correlated with the number of activated promoting regions. (2) binarized gene counts are shown to also have enough ability in distinguishing cell types^55^.

### 4.3 Post-processing procedure

After training the model, we calculate a pairwise distance matrix between cells from all batches using cell factor values. We then construct a neighborhood graph from the distance matrix by connecting each cell with both its within-batch nearest neighbors and its cross-batches nearest neighbors. Denoting the overall number of nearest neighbors for each cell by *k* (*k* = 30 for most of the results shown), the number of nearest neighbors taken in each batch is proportional to the total number of cells in the batch. More specifically, the number of neighbors *k*_*i*_ for batch *i* can be calculated by *k*_*i*_ = (*N*_*i*_*/N*_total_) · *k*, where *N*_*i*_ is the number of cells in batch *i*, and *N*_total_ is the total number of cells in all batches. We also offer an option to prune the connections in the neighborhood graph using a radius parameter *r*. The radius parameter is from 0 to 1, denoting the percentage of connections to be preserved between every two batches.

After obtaining the neighborhood graph, we then normalize the distances between cells in the graph. We first calculate the *mean within-batch distance* and *mean cross-batches distances* for each cell using the distance of the cells to its within-batch nearest neighbors and cross-batches nearest neighbors. Then we normalize the distances between the cell and its cross-batches nearest neighbors, which makes the *mean within-batch distance* and *mean cross-batches distances* for the cell to be the same. Considering cell *m* and cell *n* are nearest neighbor calculated from batch *i* and batch *j*, the distance *d*_*mn*_ between *m* and *n* can be normalized by 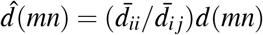, where 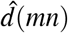 is the normalized distance between cell *m* and cell *n*, 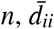 is the *mean within-batch distance* of cell *m* and its neighbors in batch *i*, 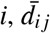 is the *mean cross-batches distance* of cell *m* and its neighbors in batch *j*. The normalized neighborhood graph can be used for visualization and clustering purposes. UMAP can take the neighborhood graph to visualize the cell-to-cell variation, and Leiden clustering algorithm is used to cluster the cells based on the neighborhood graph.

### 4.4 Retraining procedure

After clustering the cells, we use the cluster label for the retraining of scMoMaT. We first construct binary cell factor matrices from the cluster label by making each column dimension of the cell factor matrices match one specific cell cluster, and by assigning 1 to the corresponding cluster dimension and 0 to the other dimensions for each cell. The retraining step is to learn feature factors and association matrices that are consistent with the binary cell factors.

We then fix the binary cell factor matrices and update the remaining parameters in scMoMaT to minimize the loss (Eq. 2). The retrained feature factor matrices and association matrices can be used to build the *feature scoring matrices* that includes the marker score for each feature in each cell cluster. The top-scoring features in each cluster are considered to be the bio-markers of the cluster. Given the retrained feature factor matrix **C**_feat_ (e.g. **C**_*g*_, **C**_*r*_, **C**_*p*_) and shared association matrix Σ, the feature scoring matrix **M**_feat_ can be calculated as **M**_feat_ = **C**_feat_ · Σ^*T*^, and each column of **M**_feat_ are the marker scores of all features in the corresponding cell cluster.

During the retraining process, scMoMaT is flexible on the data matrices that are used for each data batch. One can incorporate additional data matrices that measure different data modalities of the existing data batches into the retraining process and learn the factor of the newly added data modalities through scMoMaT. In the testing result of mouse brain cortex dataset, PBMC dataset, BMMC dataset, we obtained the motif deviation matrices (cell by motif matrices, calculated from scATAC-seq matrix using chromVAR), and included the motif deviation matrices in the retraining process to learn the motif factor of the dataset.

### 4.5 Hyper-parameter setting in scMoMaT

There are four hyper-parameters in scMoMaT: the latent dimension *d* for cell and feature factors, the regularization weight *λ* in the loss function, the number of neighbors *k* and radius parameter *r* in the post-processing step.

The latent dimension *d* corresponds to the number of latent biological factors that should be included in the dataset. It varies according to the complexity of the cell-cell variation in the dataset. The higher the complexity is, the larger *d* is required. It does not correspond to the number of cell types and is usually larger than the number of cell types. In all our tests on real datasets, we select *d* = 30. In all our tests on simulated datasets, where the dataset structure is less complex, we set *d* = 20. The regularization weight *λ* is selected to be 0.001 (default value) for all our tests.

The number of neighbors *k* in the post-processing step should be based on the total number of cells (larger *k* for larger dataset). We suggest users to set *k* to be 30 to 50. The radius parameter *r* prunes weak connections in the neighborhood graph. Smaller *r* means more connections are removed. We suggest to apply the pruning step when there is a strong mismatch between cell type composition of different cell batches. We did not apply pruning for the real datasets; and on simulated datasets, we applied pruning and set *r* = 0.7 because of the mismatch of cell types between cell batches.

### 4.6 Data simulation

We implemented a simulation procedure which can generate multiple batches of paired scRNA-seq and scATAC-seq datasets which is generalized upon previous work SymSim^18^. Cross modality relationship between scRNA-seq and scATAC-seq data in the same batch is considered in this simulator, through the kinetic model used to generate the scRNA-seq data. More details on the simulation procedure are described in^10^. We set the number of genes to be 1000 in the scRNA-seq data matrices and the number of regions in the scATAC-seq data matrices to be 3000. 6 data batches were generated with a total number of 10000 cells.

### 4.7 Pre-processing of datasets

When running scMoMaT on real datasets, we filter genes in the scRNA-seq matrices by selecting highly variable genes. Then we quantile normalize the scRNA-seq matrices. We quantile normalize and log-transform the protein abundance matrices^56^. No protein filtering step is conducted as there is a small number of proteins measured. The scATAC-seq is filtered by selecting the regions that lie within the 2000 base-pair upstream activation region and the gene body of all genes kept in the scRNA-seq count matrices. When dealing with multiple scATAC-seq matrices with different region features, we remap the fragment file of other scATAC-seq matrices using the peaks from one scATAC-seq matrix that we select, which was also used in Cobolt^57^ and Signac^58^. This allows the scATAC-seq matrices to have the same region features.

With simulated datasets, we did not filter the genes or regions for all integration methods. The pseudo-scRNA-seq matrices are calculated by multiplying the “region by gene” association matrix that is provided by the simulator with the scATAC-seq matrices. We quantile normalize the scRNA-seq matrix, and binarize the scATAC-seq matrix when inputting these data to scMoMaT. When running UINMF and MultiMAP on the simulated data, we followed the online tutorial of these methods (see Sec. 4.8).

Some details in the preprocessing procedures can vary for each real dataset, which are described as following.

#### 4.7.1 Human PBMC dataset

For the human PBMC dataset, we selected top 7000 highly variable genes using *scanpy* for each scRNA-seq matrix separately. We do not remap the scATAC-seq matrix as the dataset comes with the same region features in the two scATAC-seq matrices. After gene and region filtering, we obtained overlapping 4768 genes, 17442 regions and 216 proteins.

#### 4.7.2 Mouse brain cortex dataset

Since the scATAC-seq matrices have different sets of region features, we first remapped the scATAC-seq matrices in the first and the fifth batches to the regions in the third batch. We then selected the top 2000 highly variable genes using *scanpy* for the scRNA-seq matrix in the second batch, and used the same set of genes for all the scRNA-seq matrices. We filled in pseudo-scRNA-seq matrix for the batches without scRNA-seq matrices, and selected the regions in scATAC-seq matrices that lie within the 2000 base-pair upstream or the gene body of the genes in scRNA-seq matrices. After the filtering process, we obtained overlapping 1677 genes and 25734 regions for all data matrices.

We download the cell label from the original data manuscripts, reorganize the labels to make them as consistent as possible. We re-annotate the “E2Rasgrf2”, “E3Rmst” and “E3Rorb” as “L2/3”, “E4Il1rapl2”, “E4Thsd7a”, “E5Galnt14”, “E5Parm1”, “E5Sulf1”, and “E5Tshz2” as “L4/5”, “E6Tle4” as “L6”, “OliM” and “OliI” as “Oligo”, “InV” as “CGE”, “InS” as “Sst”, “InP” as “Pvalb”, “InN” as “Npy”, and “Mic” as “MGC” in the first batch. We re-annotate “Lamp5”, “Vip” and “Sncg” as “CGE”, “L4”, “L5 ET” and “L5 IT” as “L4/5”, “L6 CT”, “L6 IT” and “L6b” as “L6”, “L5/6 NP” as “NP”, “Macrophage” as “MGC” in the second batch. We re-annotate “L5.IT.a”, “L5.IT.b” and “L4” as “L4/5”, “L6.CT” and “L6.IT” as “L6”, “L23.a”, “L23.b”, and “L23.c” as “L2/3”, “OGC” as “Oligo”, “ASC” as “Astro”, and “Pv” as “Pvalb” in the third batch. We re-annotate “L2/3 IT” as “L2/3”, “L4”, “L5 IT”, and “L5 PT” as “L4/5”, “L6 CT”, “L6 IT”, and “L6b” as “L6”, “Macrophage” as “MGC”, “Lamp5”, “Vip”, and “Sncg” as “CGE” in the fourth and the fifth batches.

#### 4.7.3 Healthy human BMMC dataset

We selected top 1000 highly variable genes using *scanpy* for scRNA-seq matrix. We also remove the genes with no regions in the scATAC-seq matrices lying within the 2000 bp of their upstream sequence. The filtering process gives us 924 genes 22133 regions.

#### 4.7.4 Mouse spleen dataset

We selected top 3000 highly variable genes using *scanpy* for the scRNA-seq matrix. We also remove the genes with no regions in the scATAC-seq matrices lying within the 2000 bp of their upstream sequence. The filtering process gives us 2708 genes 20435 regions.

### 4.8 Running baseline methods

#### 4.8.1 UINMF

We followed its online tutorial (http://htmlpreview.github.io/?https://github.com/welch-lab/liger/blob/master/vignettes/SNAREseq_walkthrough.html) to run UINMF. When applying UINMF to real datasets, we first binned the genome into bins of 100,000 bp for raw scATAC-seq matrices (as described in the tutorial). We then used the same the pseudo-scRNA-seq and scRNA-seq matrices as the ones that were used in scMoMaT, but normalized the matrices following UINMF tutorial instead of performing quantile normalization and log transform. We select the latent dimension to be 30 as was recommended in the tutorial. When running on simulated dataset, We generated pseudo-scRNA-seq matrices following the same procedure as scMoMaT. Setting the number of latent dimension to 30 led to very bad results so we used the ground truth number of clusters as the number of latent dimensions.

#### 4.8.2 MultiMap

We ran MultiMap following the example in its GitHub repository (https://github.com/Teichlab/MultiMAP). We ran the method using the raw data matrices as was required by the example and generated the pseudo-scRNA-seq matrices following the same procedure as was used in scMoMaT. MultiMap can be directly applied to integration scenarios with no batch that has paired data, including the human BMMC dataset, mouse spleen dataset, and the first scenario of the simulated dataset, following the example. Where there exist batches with more than one modality profiled, including the human PBMC dataset and the second scenario of the simulated dataset, we concatenated the count matrices for each batch and calculated the low dimensional representation using PCA on the concatenated matrices (as suggested by the authors). We ran MultiMap using the default hyper-parameter setting in its example.

### 4.9 Evaluation metrics

#### Graph Connectivity

Graph connectivity (GC) score measures how well the cells of the same cell type between batches are mixed in the latent space^22^. GC score is calculated by first constructing a kNN graph using cells from all batches. Then for each cell type, we select the cells that belong to the cell type and denote the corresponding subgraph by *G*_*c*_(*N*_*c*_, *E*_*c*_) where *c* denotes the cell type. The GC score of this cell type can be calculated as |LCC(*G*_*c*_)|*/*|*N*_*c*_|, where |LCC(*G*_*c*_)| denotes the largest number of connected cells within the subgraph *G*_*c*_, and |*N*_*c*_| denotes the total number of cells in the subgraph. The GC score of the whole dataset is the average GC score of all cell types.

#### Adjusted Rand Index (ARI)

The ARI score measures how well cells from different cell types can be correctly clustered regardless of batches using the latent embedding. After clustering the cells using the cell latent embedding obtained from different integration methods, we calculate the Adjusted Rand Index^59^ by comparing it with the ground truth cell label. Leiden clustering algorithm has one resolution parameter that decides the number of clusters. For each method, we ran Leiden clustering with different resolution parameters (from 0.1 to 10 with stepsize 0.5) and report the highest ARI score for all resolution parameters as the final result.

#### Normalized Mutual Information (NMI) Score

Similar to ARI score, NMI score also measures how well cells from different cell types can be correctly clustered using the latent embedding. NMI is calculated with both the cluster label and ground-truth label. For each method, we obtained the cluster label using Leiden clustering algorithm, ran the clustering algorithm with different resolution parameters (from 0.1 to 10 with stepsize 0.5) and report the highest NMI score for all resolution parameters.

#### k-Nearest Neighbor (kNN) Agreement Score

Given the cell type labels for a set of cells, the kNN agreement score measures the label agreement between each cell and its nearest neighbors. Intuitively, with high-quality labels, cells with different labels should be separated, so most of the cells should have the same label as their neighbors, unless the cell is at the boundary of two or more closely located clusters. Taking a scRNA-seq matrix as an example: we first construct a kNN graph of cells using pairwise distances obtained after performing PCA on the original data matrix. For each cell, we calculate the proportion of cells that share the same label with this cell in its *k* nearest neighbors, and the kNN agreement score of a dataset is the average of this proportion over all cells.

### 4.10 Data and code availability

The human PBMC dataset is available at Gene Expression Omnibus under accession number GSE156478. The first batch in the mouse brain cortex dataset can be accessed at Gene Expression Omnibus under accession number GSE126074. The second and the third batches can be accessed at NeMO Archive with accession number nemo:dat-ch1nqb7. The fourth batch can be downloaded with the link http://celltypes.brain-map.org/api/v2/well_known_file_download/694413985. The fifth batch can be downloaded from 10x Genomics website with the link https://support.10xgenomics.com/single-cell-atac/datasets/1.1.0/atac_v1_adult_brain_fresh_5k. The healthy human BMMC dataset is available at Gene Expression Omnibus under accession number GSE139369. The scATAC-seq and scRNA-seq matrix of mouse spleen dataset are available at ArrayExpress under accession numbers E-MTAB-6714 and E-MTAB-9769.

The code of scMoMaT is available at https://github.com/PeterZZQ/scMoMaT.

## Supplementary Figures and tables

**Figure S1.**
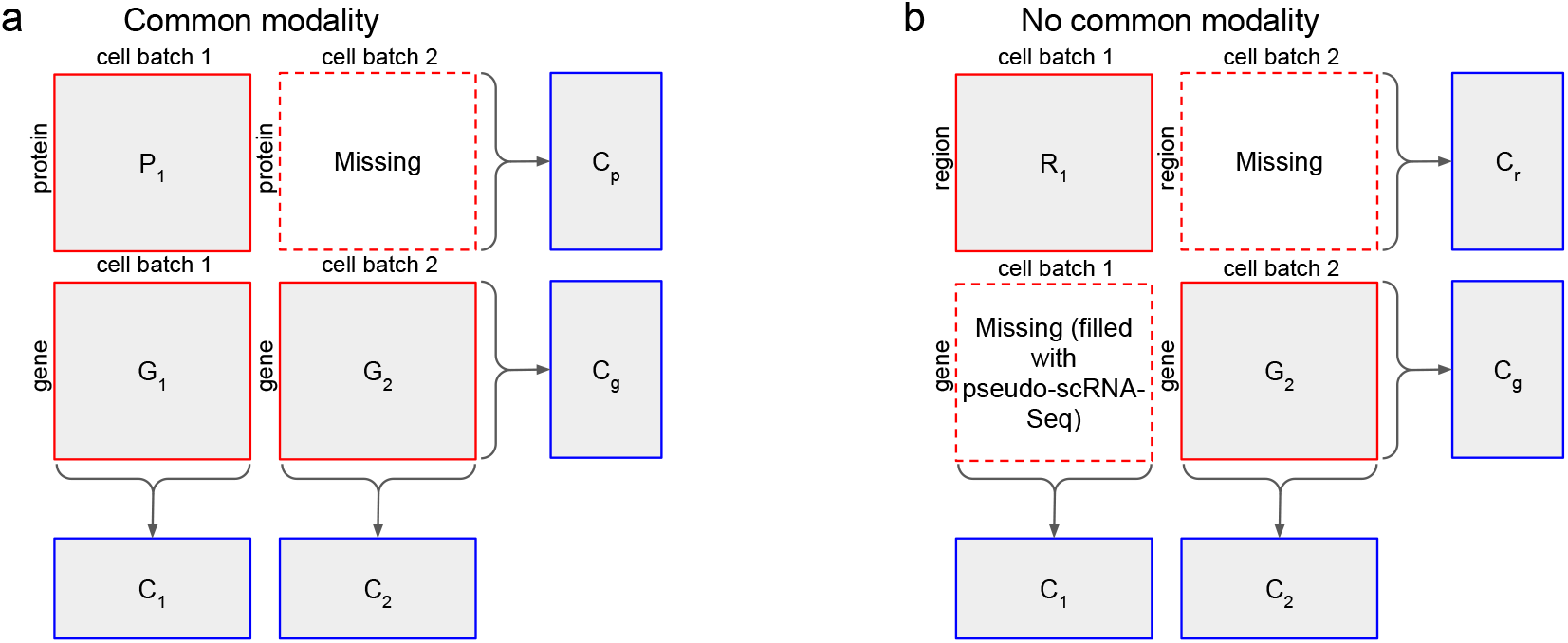
Two example integration scenarios of scMoMaT. **a**. An example where data batches have common modality. **b**. An example where data batches do not have common modality. scMoMaT fills in the missing modality using a pseudo-scRNA-seq matrix, and jointly factorizes the data matrices along with pseudo-scRNA-seq matrix.

**Figure S2.**
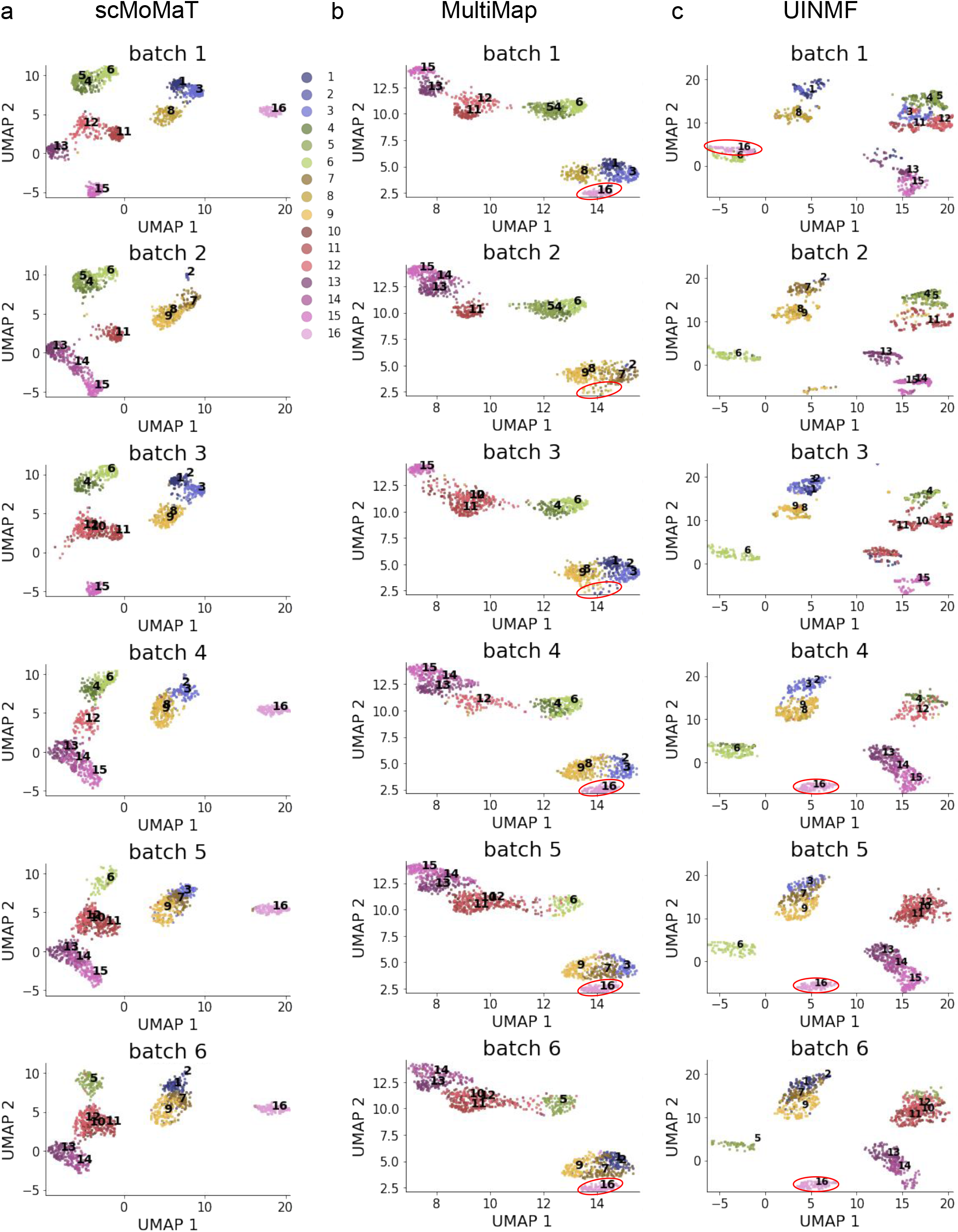
Cell embedding of scMoMaT, MultiMap, and UINMF on one example simulated dataset, visualized using UMAP. For each method, cells are plot separately for each batch, and colored by ground truth cell type labels. **a**. The cell embedding of scMoMaT; **b**. The cell embedding of MultiMap; **c**. The cell embedding of UINMF.

**Figure S3.**
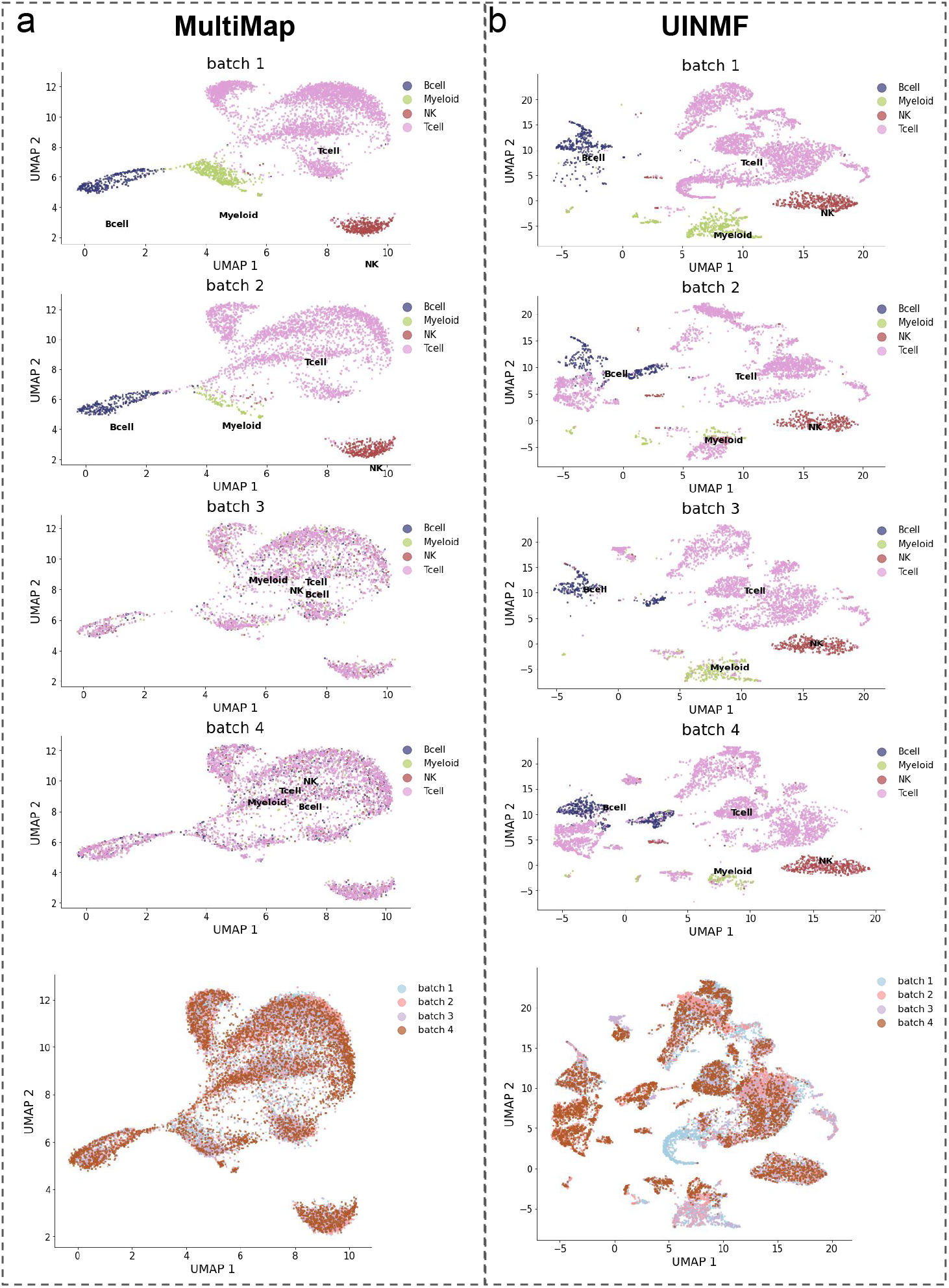
Cell embedding of MultiMap and UINMF on the human PBMC dataset, visualized using UMAP. **a**. (Upper four plots) The cell embedding of MultiMap, where cells are plot separately for different batches, and colored by cell type labels from original data paper. (Lower plot) The cell embedding of MultiMap, where cells are colored by data batches. **b**. (Upper four plots) The cell embedding of UINMF, where cells are plot separately for different batches, and colored by cell type labels from original data paper. (Lower plot) The cell embedding of UINMF, where cells are colored by data batches.

**Figure S4.**
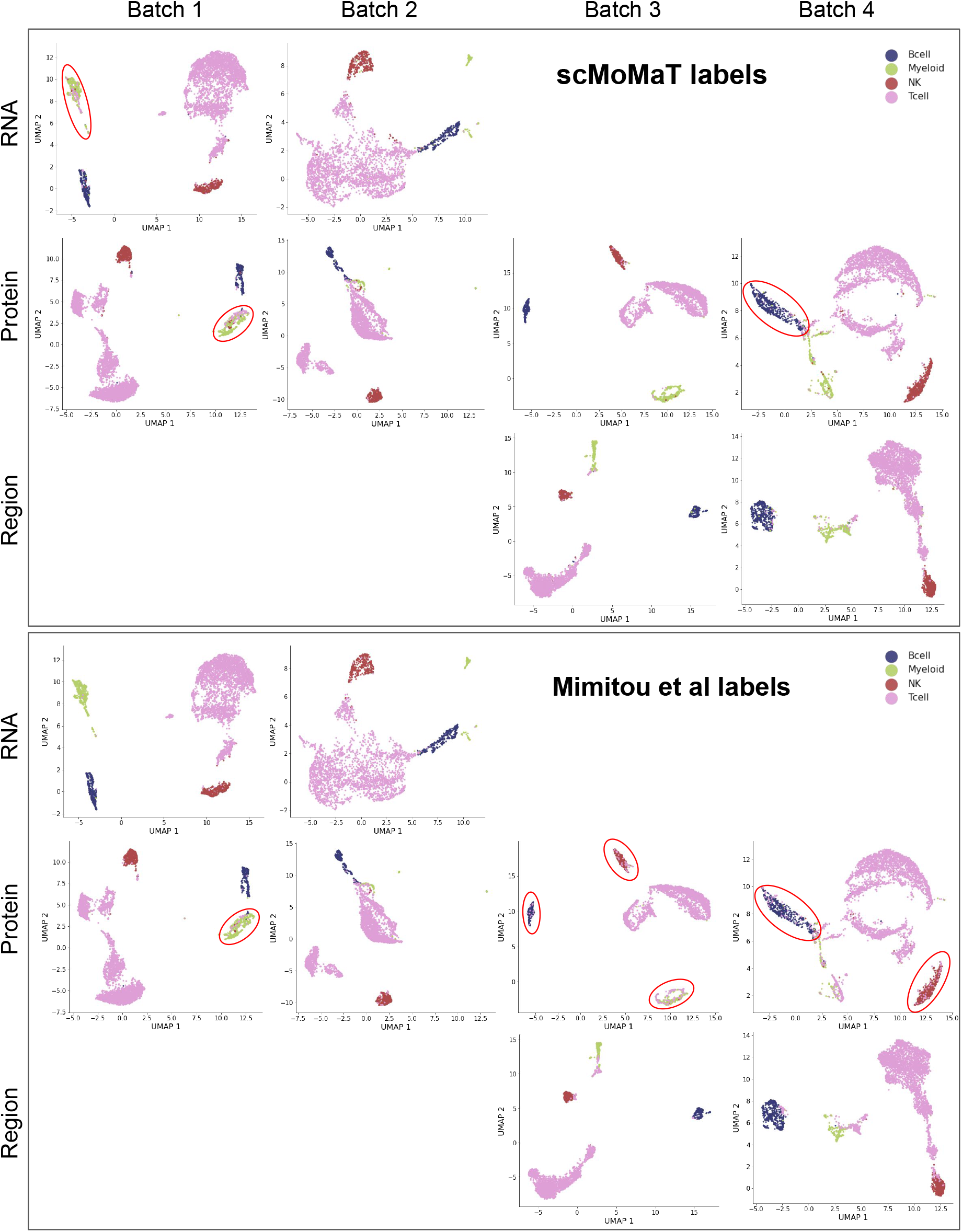
UMAP visualization of each of the 8 data matrices in human PBMC dataset, where cells are colored by the cell type label from scMoMaT, and the cell type label in the original data paper (Mimitou et al).

**Figure S5.**
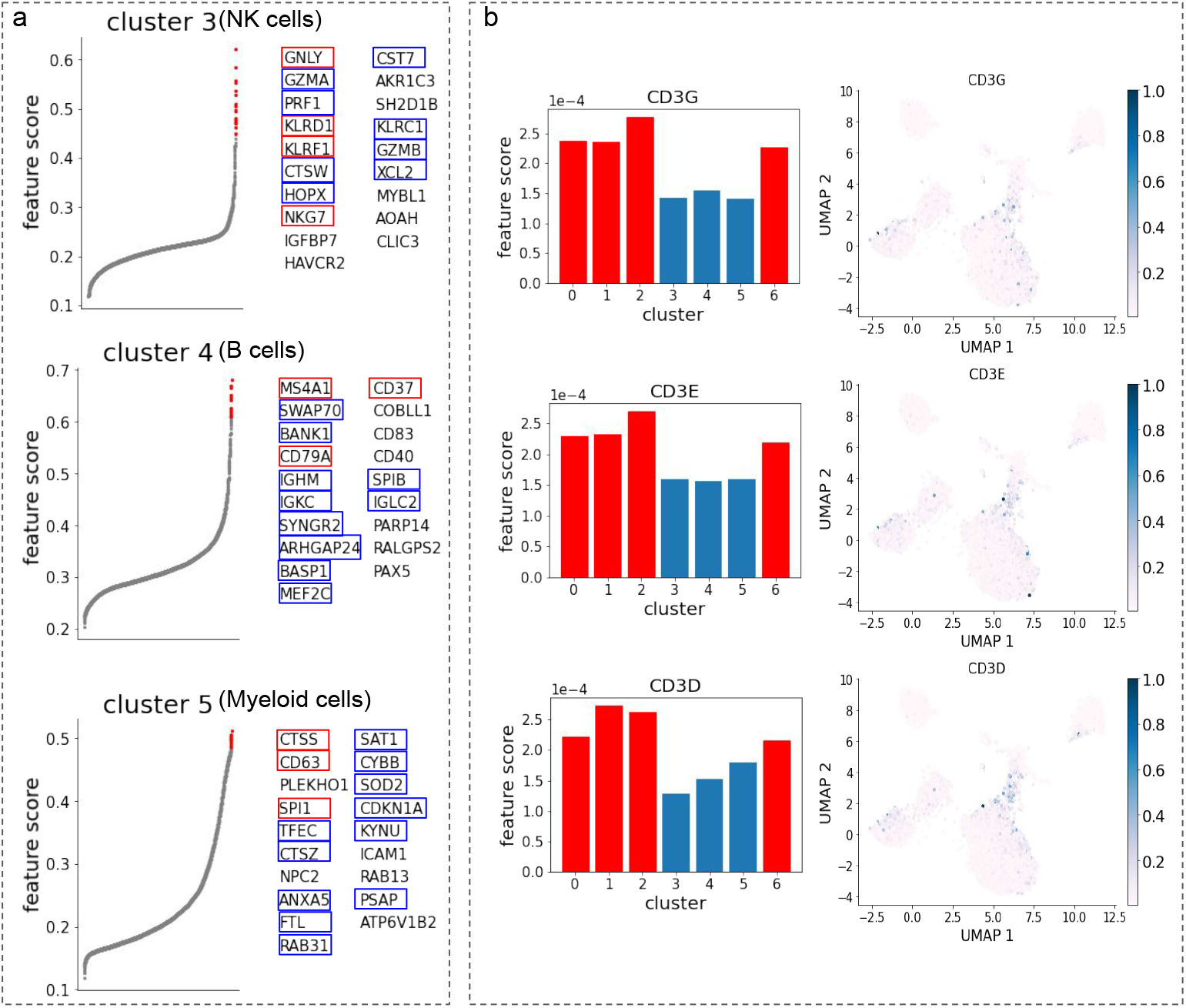
Additional results on human PBMC dataset. **a**. The top-20 scoring genes in cluster 3 (NK cells), 4 (B cells), and 5 (Myeloid cells). Known marker genes are annotated in red and blue frames. **b**. (Left) The scores of T cell marker genes *CD3G, CD3E*, and *CD3D* of different Leiden clusters, where x-axis corresponds to Leiden clusters. Top-scoring clusters are colored red. (Right) Abundance levels of *CD3G, CD3E*, and *CD3D* on the cell embedding of scMoMaT.

**Figure S6.**
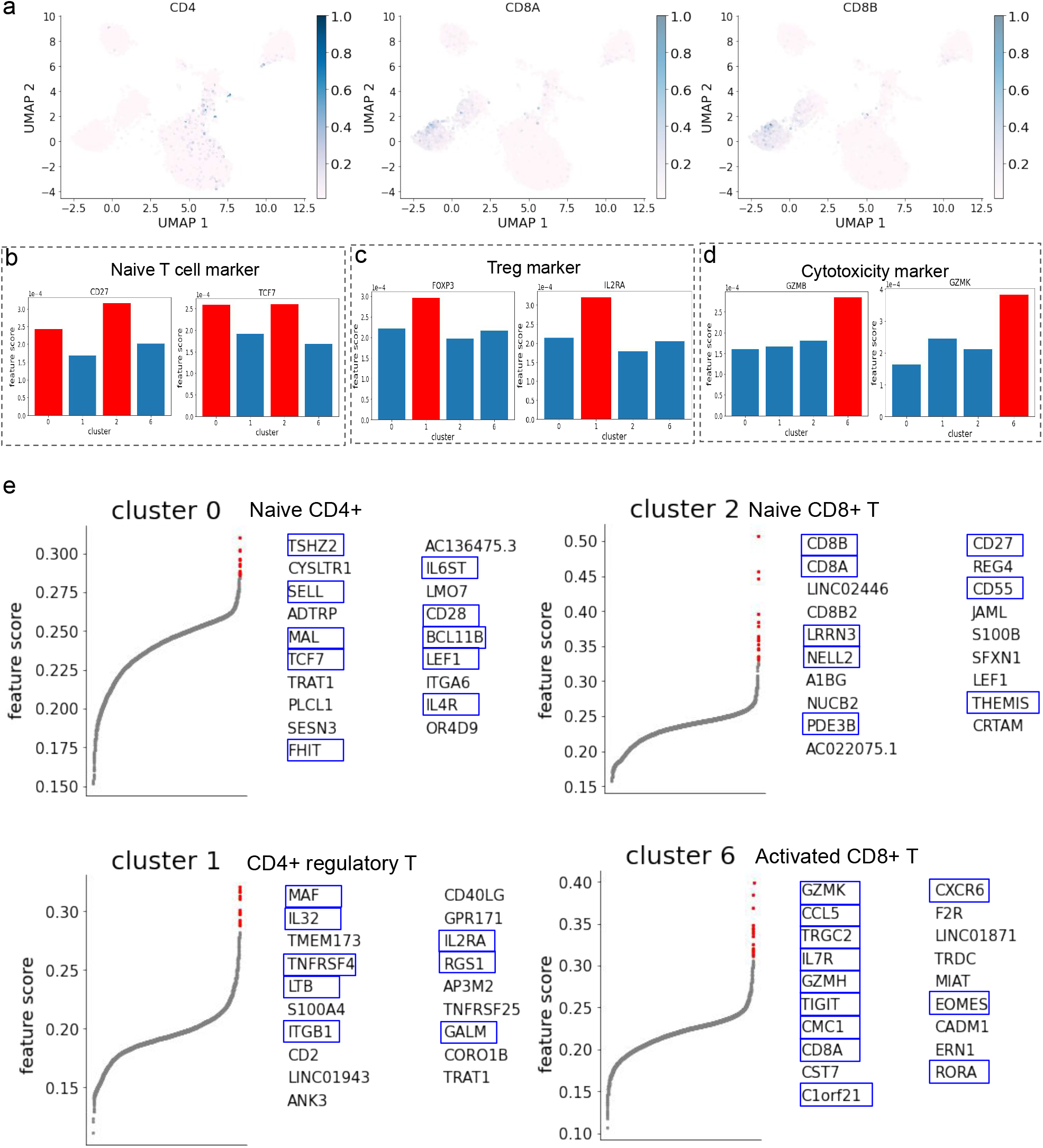
Additional results on human PBMC dataset. **a**. Abundance levels of *CD4, CD8A*, and *CD8B* on the cell embedding of scMoMaT. **b-d**. scores of (**b**) Navie T cell marker genes, (**c**) Treg cell marker genes, and (**d**) Cytotoxicity cell marker genes in different Leiden clusters, where x-axis corresponds to Leiden clusters. **e**. The top-20 scoring genes of cluster 0 (Naive CD4+), 2 (Naive CD8+), 1 (CD4+ regulatory), and 6 (activated CD8+) T cells. Known marker genes from literature are annotated in blue frames.

**Figure S7.**
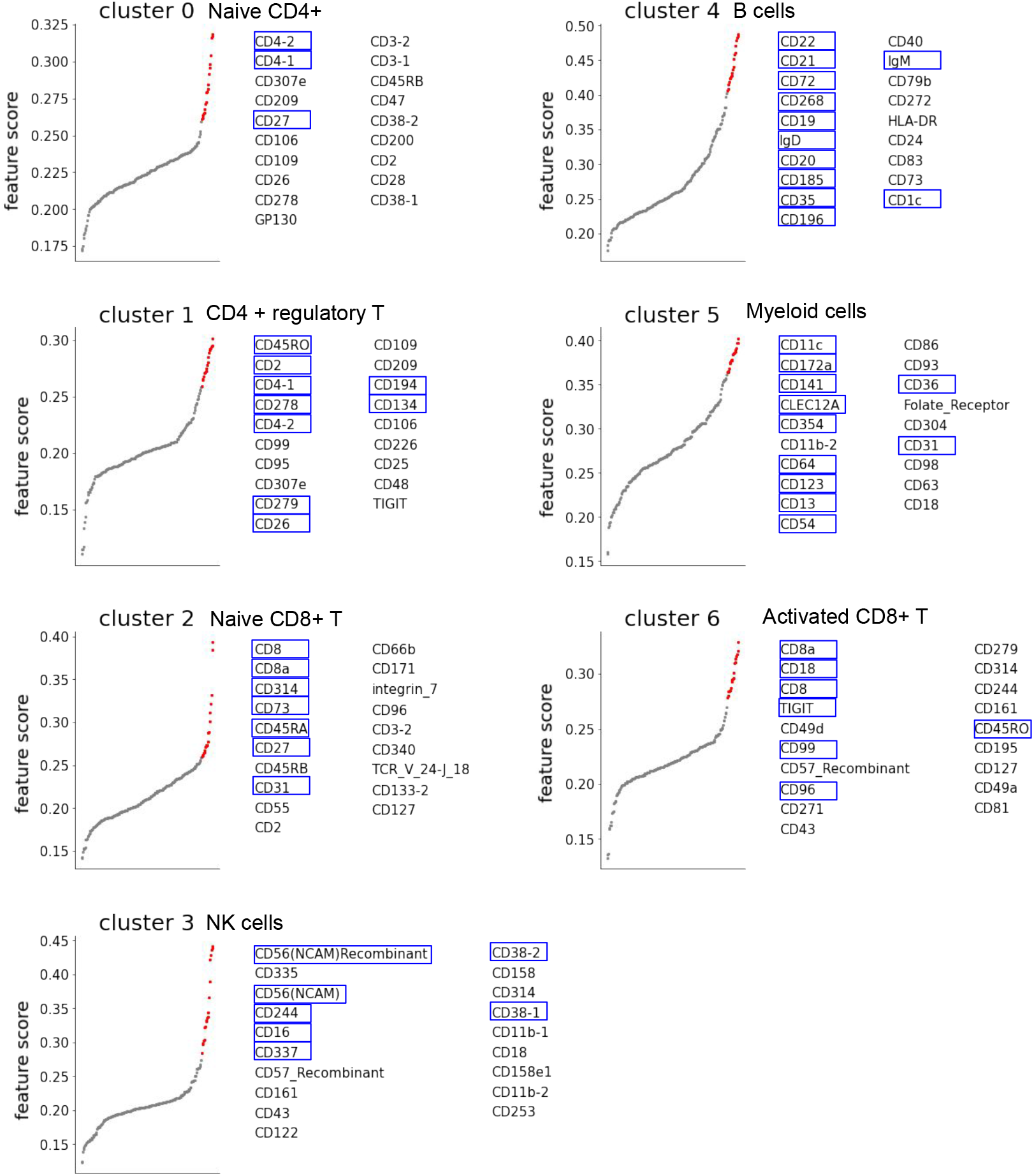
Top-20 scoring proteins in all Leiden clusters of human PBMC dataset. Known marker proteins are annotated in blue frames.

**Figure S8.**
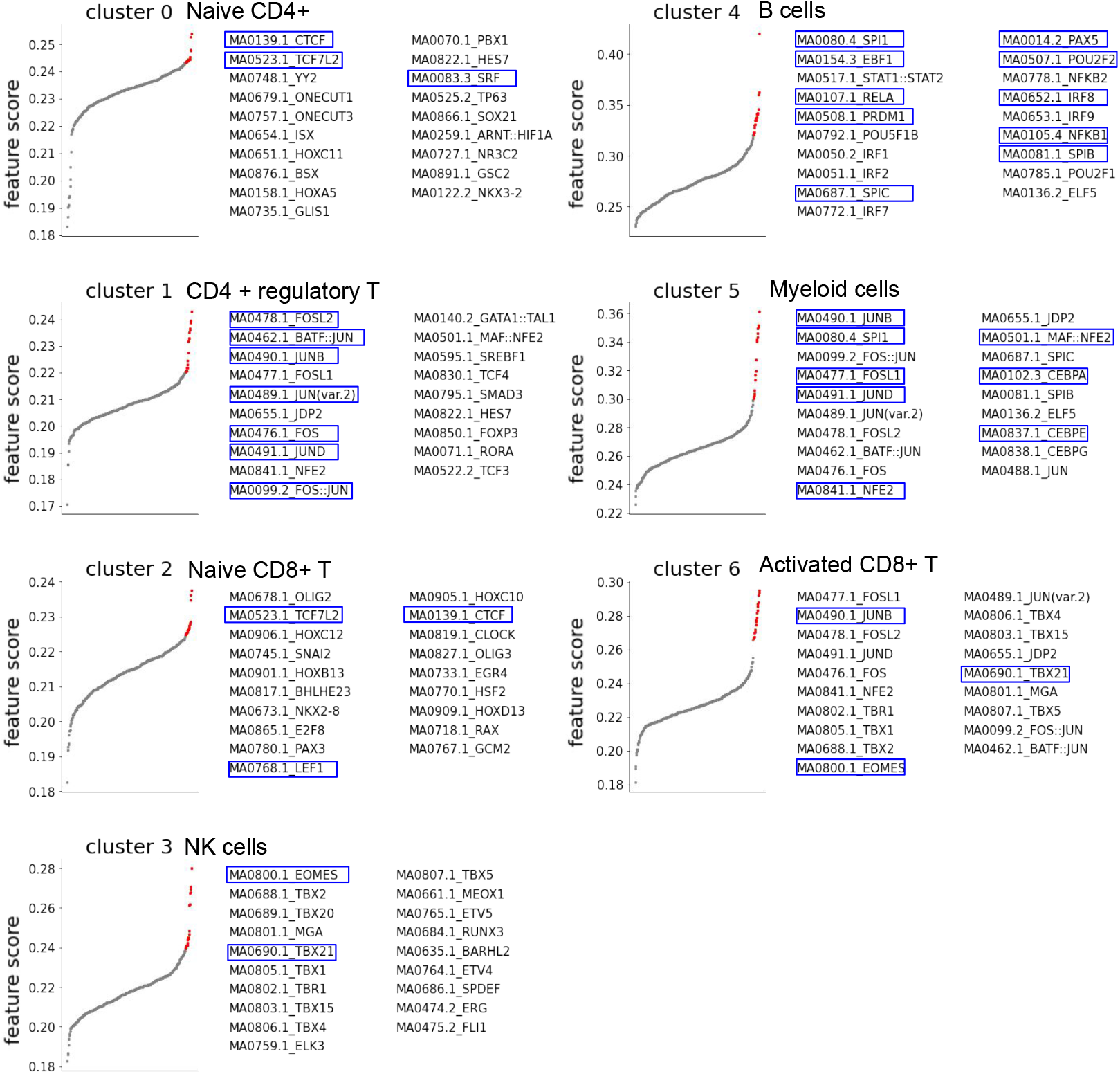
Top-20 scoring motifs in all Leiden clusters of human PBMC dataset. Marker motifs corresponding to know TFs for each cell type are annotated in blue frames.

**Figure S9.**
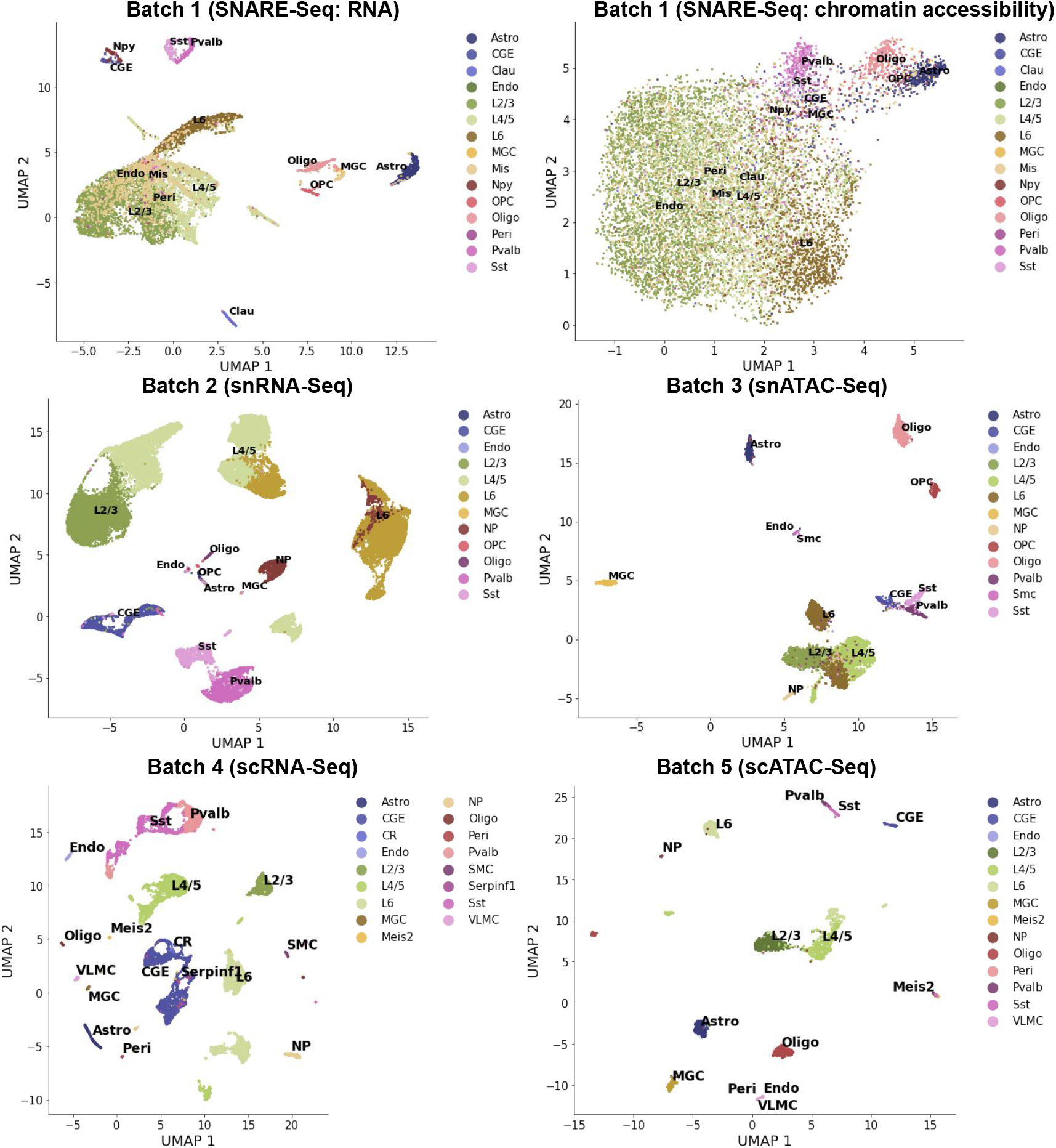
UMAP visualization of 6 data matrices in mouse brain cortex dataset. Cells are colored by the cell types re-organized from the original data paper.

**Figure S10.**
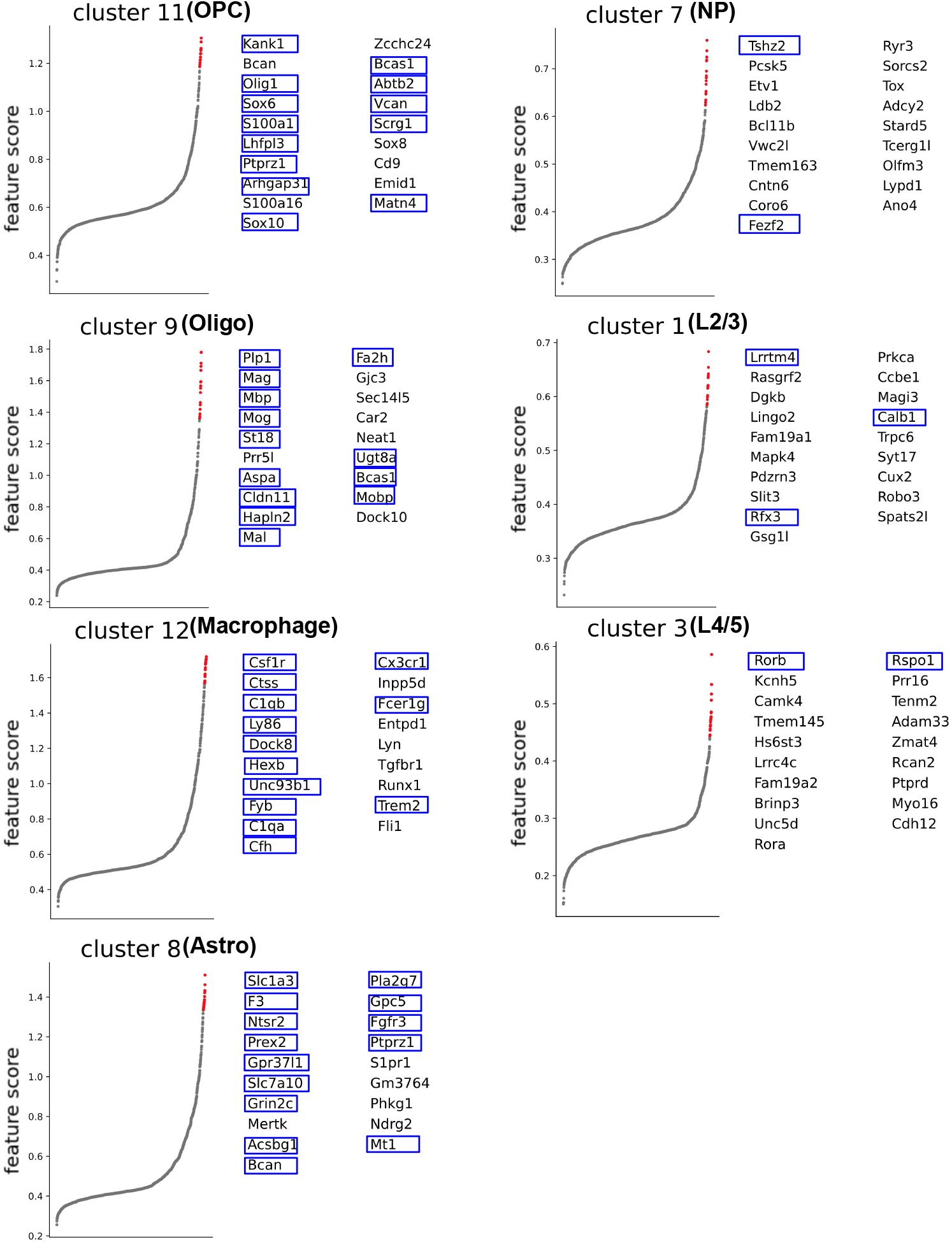
Top-20 scoring genes in Leiden clusters 11 (OPC), 9 (Oligo), 12 (Macrophage), 8 (Astro), 7 (NP), 1 (L2/3), 3 (L4/5) of mouse brain cortex dataset. Known marker genes are annotated in blue frames.

**Figure S11.**
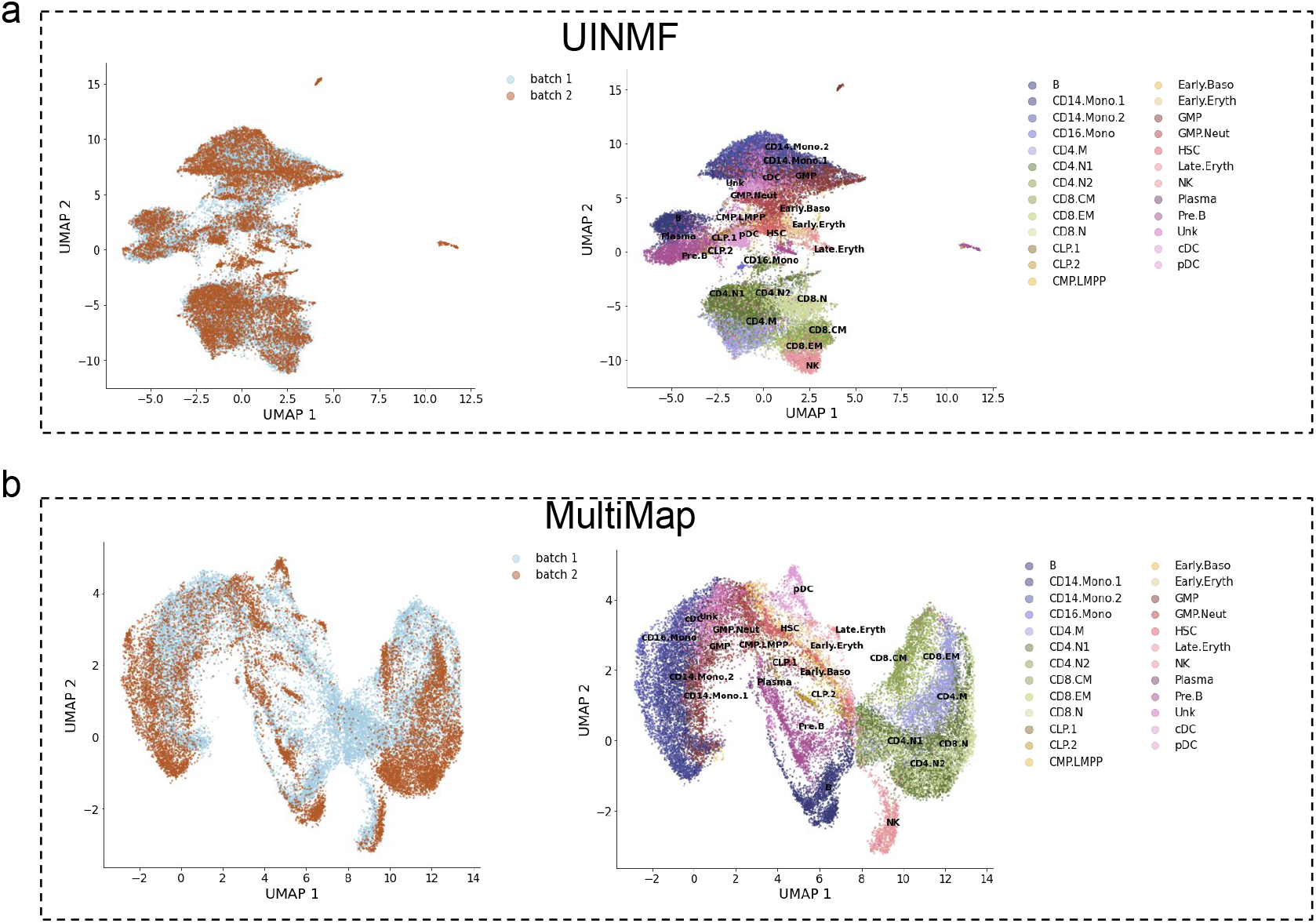
Cell embedding of UINMF and MultiMap for human bone marrow dataset. **a**. Cell embedding of UINMF for human bone marrow dataset, where cells are colored by (left) data batches, and (right) cell type labels from original data paper. **b**. Cell embedding of MultiMap for human bone marrow dataset, where cells are colored by (left) data batches, and (right) cell type labels from original data paper.

**Figure S12.**
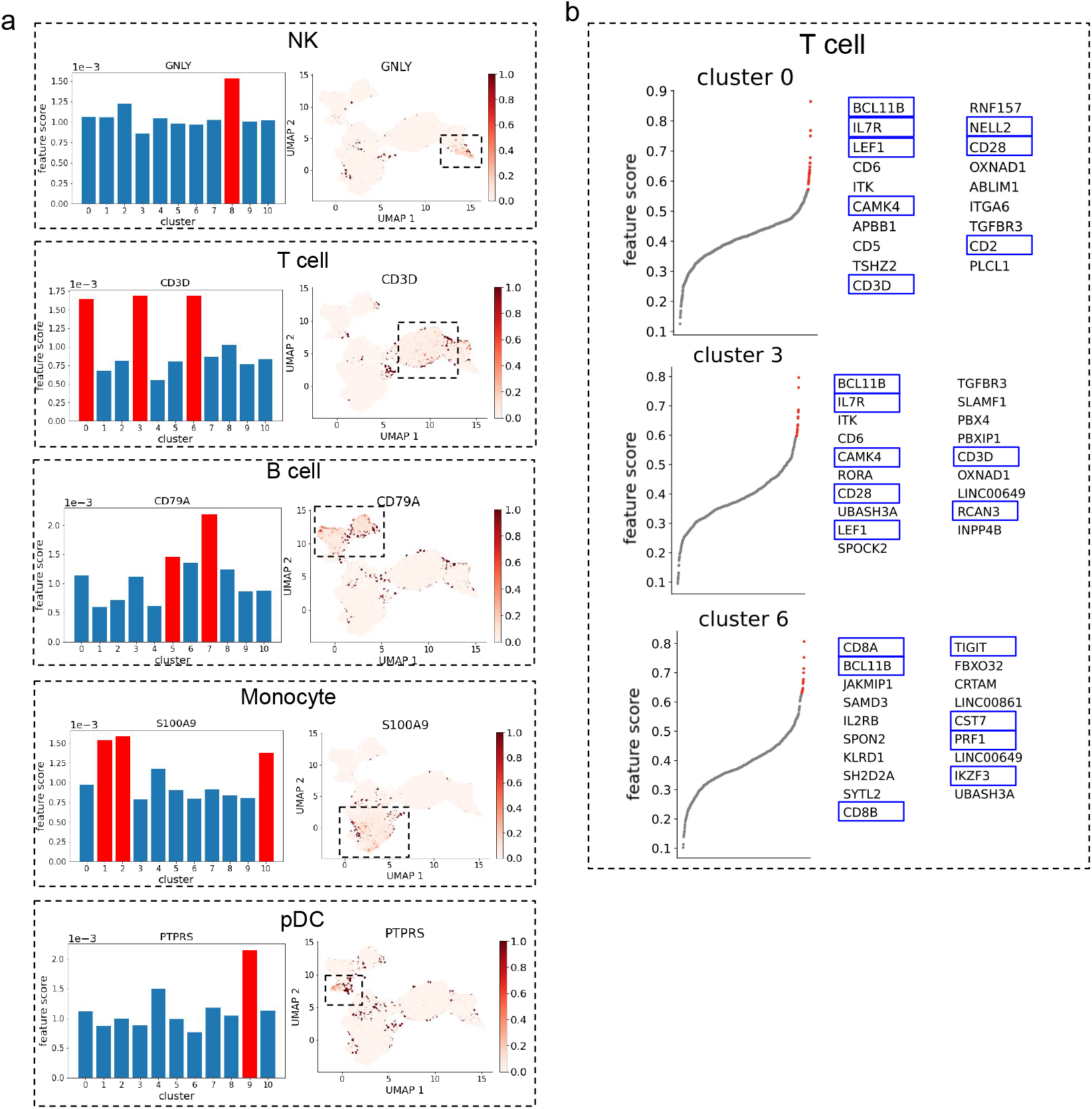
Results on the gene factors learned from human bone marrow dataset. **a**. (Left) The scores of NK, T, B, Monocyte and pDC cell marker genes in different Leiden clusters, where x-axis correspond to Leiden clusters. (Right) Abundance level of these marker genes on cell embedding learned from scMoMaT. **b**. The top-20 scoring genes of the Leiden clusters that correspond to T cell (cluster 0, 3, and 6). Known marker genes are annotated in blue frames.

**Figure S13.**
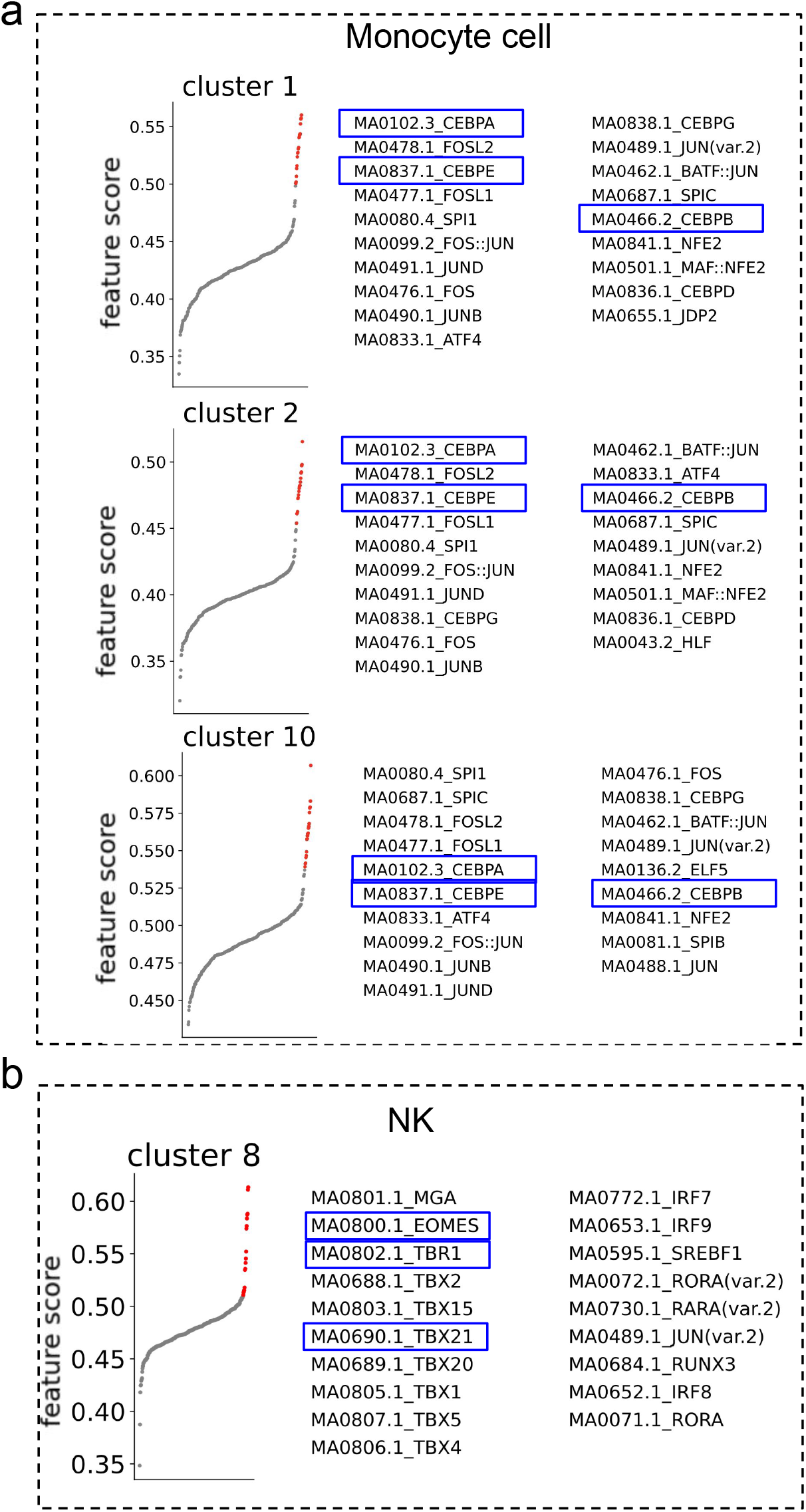
The motif factors learned from human bone marrow dataset. **a**. The top-20 scoring motifs of the Leiden clusters that correspond to Monocyte cells (cluster 1, 2, and 10), where known marker motifs are annotated in blue frames. **b**. The top-20 scoring motifs of the Leiden cluster that correspond to NK cells (cluster 8), where known marker motifs are annotated in blue frames.

**Figure S14.**
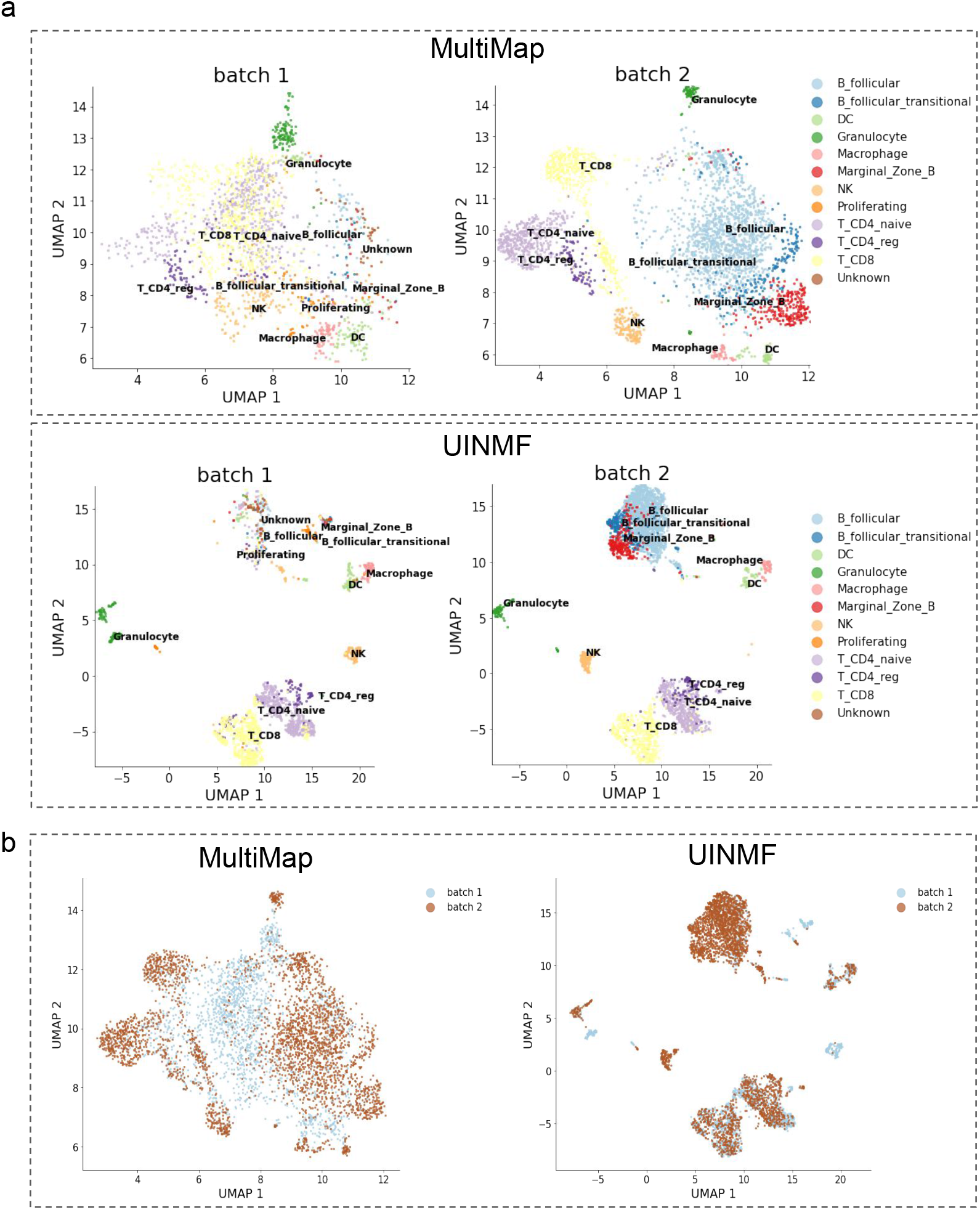
Cell embedding of MultiMap and UINMF for mouse spleen dataset (after sub-sampling). Cells are colored by (**a**) cell type annotation in the original data paper and (**b**) data batches.

**Table S2.**
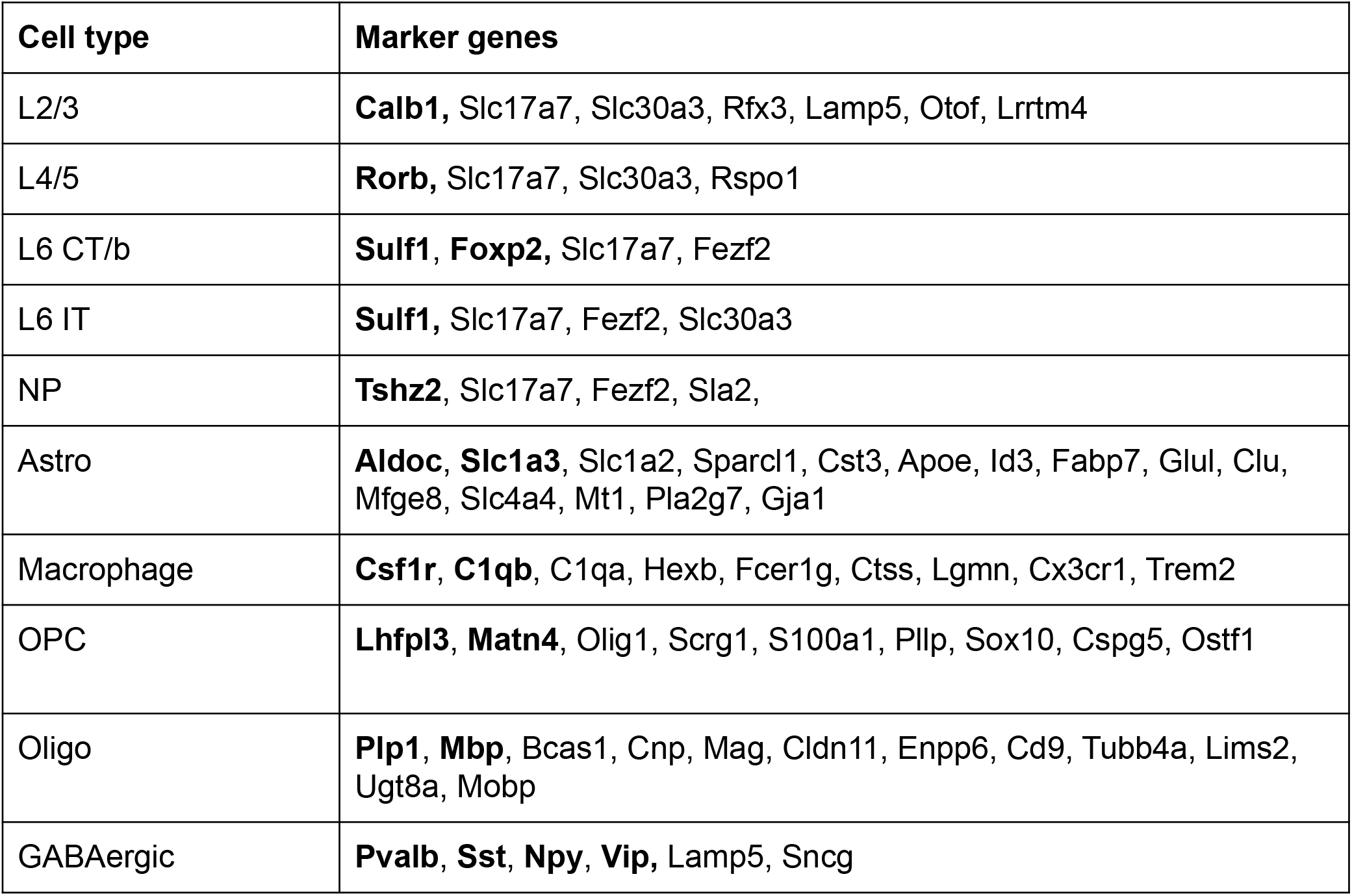
Marker gene list of different cell types in mouse brain cortex dataset, collected from literature.

## References

1. Stoeckius, M. et al. Simultaneous epitope and transcriptome measurement in single cells. Nat. Methods 14, 865–868 (2017).

2. Chen, S., Lake, B. B. & Zhang, K. High-throughput sequencing of the transcriptome and chromatin accessibility in the same cell. Nat. biotechnology 37, 1452–1457 (2019).

3. Mimitou, E. P. et al. Scalable, multimodal profiling of chromatin accessibility, gene expression and protein levels in single cells. Nat. Biotechnol. (2021).

4. Argelaguet, R., Cuomo, A. S. E., Stegle, O. & Marioni, J. C. Computational principles and challenges in single-cell data integration. Nat. Biotechnol. 1–14 (2021).

5. Welch, J. D. et al. Single-Cell multi-omic integration compares and contrasts features of brain cell identity. Cell 177, 1873–1887.e17 (2019).

6. Stuart, T. et al. Comprehensive integration of Single-Cell data. Cell 177, 1888–1902.e21 (2019).

7. Duren, Z. et al. Integrative analysis of single-cell genomics data by coupled nonnegative matrix factorizations. Proc. Natl. Acad. Sci. U. S. A. 115, 7723–7728 (2018).

8. Cao, K., Bai, X., Hong, Y. & Wan, L. Unsupervised topological alignment for single-cell multi-omics integration. Bioinformatics 36, i48–i56 (2020).

9. Singh, R. et al. Unsupervised manifold alignment for single-cell multi-omics data. In Proceedings of the 11th ACM International Conference on Bioinformatics, Computational Biology and Health Informatics, 1–10 (Association for Computing Machinery, New York, NY, USA, 2020).

10. Zhang, Z., Yang, C. & Zhang, X. scDART: integrating unmatched scRNA-seq and scATAC-seq data and learning cross-modality relationship simultaneously. Genome Biol. 23, 139 (2022).

11. Hao, Y. et al. Integrated analysis of multimodal single-cell data. Cell (2021).

12. Jin, S., Zhang, L. & Nie, Q. scAI: an unsupervised approach for the integrative analysis of parallel single-cell transcriptomic and epigenomic profiles. Genome Biol. 21, 25 (2020).

13. Ashuach, T., Gabitto, M. I., Jordan, M. I. & Yosef, N. Multivi: deep generative model for the integration of multi-modal data. bioRxiv (2021).

14. Jain, M. S. et al. MultiMAP: dimensionality reduction and integration of multimodal data. Genome Biol. 22, 346 (2021).

15. Hao, Y. et al. Dictionary learning for integrative, multimodal, and scalable single-cell analysis. bioRxiv (2022).

16. Kriebel, A. R. & Welch, J. D. Uinmf performs mosaic integration of single-cell multi-omic datasets using nonnegative matrix factorization. Nat. communications 13, 1–17 (2022).

17. Schep, A. N., Wu, B., Buenrostro, J. D. & Greenleaf, W. J. chromvar: inferring transcription-factor-associated accessibility from single-cell epigenomic data. Nat. methods 14, 975–978 (2017).

18. Zhang, X., Xu, C. & Yosef, N. Simulating multiple faceted variability in single cell RNA sequencing. Nat. Commun. 10, 2611 (2019).

19. Lee, D. D. & Seung, H. S. Learning the parts of objects by non-negative matrix factorization. Nature 401, 788–791 (1999).

20. Traag, V. A., Waltman, L. & Van Eck, N. J. From louvain to leiden: guaranteeing well-connected communities. Sci. reports 9, 1–12 (2019).

21. Jain, M. S. et al. Multimap: dimensionality reduction and integration of multimodal data. Genome biology 22, 1–26 (2021).

22. Luecken, M. D. et al. Benchmarking atlas-level data integration in single-cell genomics. BioRxiv (2020).

23. Stoeckius, M. et al. Simultaneous epitope and transcriptome measurement in single cells. Nat. methods 14, 865–868 (2017).

24. Mimitou, E. P. et al. Multiplexed detection of proteins, transcriptomes, clonotypes and crispr perturbations in single cells. Nat. methods 16, 409–412 (2019).

25. Yang, C. et al. Heterogeneity of human bone marrow and blood natural killer cells defined by single-cell transcriptome. Nat. communications 10, 1–16 (2019).

26. Stelzer, G. et al. The GeneCards suite: From gene data mining to disease genome sequence analyses. Curr. Protoc. Bioinforma. 54, 1.30.1–1.30.33 (2016).

27. Xu-Monette, Z. Y. et al. Assessment of cd37 b-cell antigen and cell of origin significantly improves risk prediction in diffuse large b-cell lymphoma. Blood, The J. Am. Soc. Hematol. 128, 3083–3100 (2016).

28. Tang-Huau, T.-L. et al. Human in vivo-generated monocyte-derived dendritic cells and macrophages cross-present antigens through a vacuolar pathway. Nat. communications 9, 1–12 (2018).

29. Hauses, M., Tönjes, R. R. & Grez, M. The transcription factor sp1 regulates the myeloid-specific expression of the human hematopoietic cell kinase (hck) gene through binding to two adjacent gc boxes within the hck promoter-proximal region. J. Biol. Chem. 273, 31844–31852 (1998).

30. Knol, E. F., Mul, F. P., Jansen, H., Calafat, J. & Roos, D. Monitoring human basophil activation via cd63 monoclonal antibody 435. J. Allergy Clin. Immunol. 88, 328–338 (1991).

31. Zhang, X. et al. CellMarker: a manually curated resource of cell markers in human and mouse. Nucleic Acids Res. 47, D721–D728 (2019).

32. Johannisson, A. & Festin, R. Phenotype transition of cd4+ t cells from cd45ra to cd45ro is accompanied by cell activation and proliferation. Cytom. The J. Int. Soc. for Anal. Cytol. 19, 343–352 (1995).

33. Caccamo, N., Joosten, S. A., Ottenhoff, T. H. & Dieli, F. Atypical human effector/memory cd4+ t cells with a naive-like phenotype. Front. Immunol. 2832 (2018).

34. Szabo, P. A. et al. Single-cell transcriptomics of human t cells reveals tissue and activation signatures in health and disease. Nat. communications 10, 1–16 (2019).

35. Yao, Z. et al. A transcriptomic and epigenomic cell atlas of the mouse primary motor cortex. Nature 598, 103–110 (2021).

36. Tasic, B. et al. Adult mouse cortical cell taxonomy revealed by single cell transcriptomics. Nat. neuroscience 19, 335–346 (2016).

37. Tasic, B. et al. Shared and distinct transcriptomic cell types across neocortical areas. Nature 563, 72–78 (2018).

38. Cao, Y. et al. Sailer: scalable and accurate invariant representation learning for single-cell atac-seq processing and integration. Bioinformatics 37, i317–i326 (2021).

39. Chen, Z. et al. SCAN-ATAC-Sim: a scalable and efficient method for simulating single-cell ATAC-seq data from bulk-tissue experiments. Bioinformatics 37, 1756–1758, DOI: 10.1093/bioinformatics/btaa1039 (2021). https://academic.oup.com/bioinformatics/article-pdf/37/12/1756/39119344/btaa1039.pdf.

40. Traag, V. A., Waltman, L. & van Eck, N. J. From louvain to leiden: guaranteeing well-connected communities. Sci. Rep. 9, 5233 (2019).

41. Bandler, R. C. et al. Single-cell delineation of lineage and genetic identity in the mouse brain. Nature 601, 404–409 (2022).

42. de Wit, J. et al. Unbiased discovery of glypican as a receptor for lrrtm4 in regulating excitatory synapse development. Neuron 79, 696–711 (2013).

43. Tremblay, R., Lee, S. & Rudy, B. Gabaergic interneurons in the neocortex: from cellular properties to circuits. Neuron 91, 260–292 (2016).

44. Li, Y. E. et al. An atlas of gene regulatory elements in adult mouse cerebrum. Nature 598, 129–136 (2021).

45. Mulvaney, J. & Dabdoub, A. Atoh1, an essential transcription factor in neurogenesis and intestinal and inner ear development: function, regulation, and context dependency. J. Assoc. for Res. Otolaryngol. 13, 281–293 (2012).

46. Dixit, R. et al. Neurog1 and neurog2 control two waves of neuronal differentiation in the piriform cortex. J. Neurosci. 34, 539–553 (2014).

47. Granja, J. M. et al. Single-cell multiomic analysis identifies regulatory programs in mixed-phenotype acute leukemia. Nat. biotechnology 37, 1458–1465 (2019).

48. Zhao, F. et al. S100a9 a new marker for monocytic human myeloid-derived suppressor cells. Immunology 136, 176–183 (2012).

49. Bunin, A. et al. Protein tyrosine phosphatase ptprs is an inhibitory receptor on human and murine plasmacytoid dendritic cells. Immunity 43, 277–288 (2015).

50. Marchwicka, A. & Marcinkowska, E. Regulation of expression of cebp genes by variably expressed vitamin d receptor and retinoic acid receptor α in human acute myeloid leukemia cell lines. Int. J. Mol. Sci. 19, 1918 (2018).

51. Matsushita, H. et al. C/ebpα and c/ebpε induce the monocytic differentiation of myelomonocytic cells with the mll-chimeric fusion gene. Oncogene 27, 6749–6760 (2008).

52. Kiekens, L. et al. T-bet and eomes accelerate and enhance functional differentiation of human natural killer cells. Front. immunology 12 (2021).

53. Huang, C. & Bi, J. Expression regulation and function of t-bet in nk cells. Front. Immunol. 12, 761920 (2021).

54. Chen, X., Miragaia, R. J., Natarajan, K. N. & Teichmann, S. A. A rapid and robust method for single cell chromatin accessibility profiling. Nat. communications 9, 1–9 (2018).

55. Qiu, P. Embracing the dropouts in single-cell rna-seq analysis. Nat. communications 11, 1–9 (2020).

56. Bolstad, B. M., Irizarry, R. A., Åstrand, M. & Speed, T. P. A comparison of normalization methods for high density oligonucleotide array data based on variance and bias. Bioinformatics 19, 185–193 (2003).

57. Gong, B., Zhou, Y. & Purdom, E. Cobolt: integrative analysis of multimodal single-cell sequencing data. Genome biology 22, 1–21 (2021).

58. Stuart, T., Srivastava, A., Madad, S., Lareau, C. A. & Satija, R. Single-cell chromatin state analysis with signac. Nat. methods 18, 1333–1341 (2021).

59. Steinley, D. Properties of the hubert-arable adjusted rand index. Psychol. methods 9, 386 (2004).

